# Reconstitution of surface lipoprotein translocation reveals Slam as an outer membrane translocon in Gram-negative bacteria

**DOI:** 10.1101/2021.08.22.457263

**Authors:** Minh Sang Huynh, Yogesh Hooda, Raina Li, Maciej Jagielnicki, Christine Chieh-Lin Lai, Trevor F Moraes

**Affiliations:** Department of Biochemistry, University of Toronto, Canada; MRC Laboratory of Molecular Biology, Cambridge University, UK

## Abstract

Surface lipoproteins (SLPs) are peripherally attached to the outer leaflet of the outer membrane in many Gram-negative bacteria, playing significant roles in nutrient acquisition and immune evasion in the host. While the factors that are involved in the synthesis and delivery of SLPs in the inner membrane are well characterized, the molecular machineries required for the movement of SLPs to the surface are still not fully elucidated. In this study, we investigated the translocation of a surface lipoprotein TbpB through a Slam1-dependent pathway. Using purified components, we developed an *in vitro* translocation assay where unfolded TbpB is transported through Slam1 containing proteoliposomes, confirming Slam1 as an outer membrane translocon. While looking to identify factors to increase translocation efficiency, we discovered the periplasmic chaperone Skp interacted with TbpB in the periplasm of *Escherichia coli*. The presence of Skp was found to increase the translocation efficiency of TbpB in the reconstituted translocation assays. A knockout of Skp in *Neisseria meningitidis* revealed that Skp is essential for functional translocation of TbpB to the bacterial surface. Taken together, we propose a pathway for surface destined lipoproteins, where Skp acts as a holdase for Slam-mediated TbpB translocation across the outer membrane.

## Introduction

Transport of proteins to their correct spatio-temporal location is imperative for cell survival. This key process often requires the movement of proteins across lipid bilayers through a translocation channel which is also referred to as a translocon (Walter and Lingappa, 1986; Schnell and Hebert, 2003). Translocons are found in all living organisms and include the Sec translocon that is responsible for the bulk of protein transport across the inner plasma membrane (in prokaryotes) and the endoplasmic reticulum membrane (in eukaryotes) (Johnson and Van Waes, 1999; Tsirigotaki et al, 2017). Many Gram-negative bacteria contain an additional outer membrane (OM) that is separated from the plasma membrane (or inner membrane - IM) by a periplasmic space and a peptidoglycan layer. A number of outer membrane translocons have been previously identified and use different molecular mechanisms to export proteins to the extracellular matrix (Karuppiah et al, 2011).

Surface lipoproteins (SLPs) are peripheral membrane proteins that are anchored to the surface of Gram-negative bacteria. These proteins play critical roles in bacterial physiology and virulence (Wilson and Bernstein, 2016). Many SLPs were shown to improve bacterial fitness and survival in the host environment, especially for pathogenic bacteria such as *Neisseria*, *Bacteroides* and *Spirochetes* (Hooda and Moraes, 2018). SLPs contain an N-terminal signal peptide that allows their translocation across the inner membrane by the Sec or Tat machinery. In the periplasmic space, the SLPs are modified by three biosynthetic enzymes that cleave the signal peptide and add a lipid group to the N-terminal cysteine residue which anchors them to the inner membrane (Zückert, 2014). Most of these lipidated SLPs are recognized by the Lol system, which then delivers SLPs across the periplasm to the inner leaflet of the outer membrane (Szewczyk and Collet, 2016; Okuda and Tokuda, 2011). In the outer membrane, the protein machinery responsible for the translocation of SLPs across the outer membrane is known only for a handful of SLPs. Recently, an outer membrane protein named Slam (**<u>S</u>**urface **l**ipoprotein **<u>a</u>**ssembly **<u>m</u>**odulator) was identified in the human pathogen *Neisseria meningitidis* that is necessary for the surface display of the SLP transferrin binding protein B or TbpB (Hooda et al, 2016). Slam-like proteins were subsequently found in several Gram-negative bacteria from the phylum *Proteobacteria* (Hooda et al, 2017). Co-expression of Slam1 (the first Slam discovered) with TbpB in the model organism *Escherichia coli*, which lacks both Slam1 and TbpB genes, allows for functional surface display of TbpB. Further, Slam1 was found to interact with TbpB in the outer membrane through co-immunoprecipitation experiments (Hooda et al 2016). Taken together, these results confirmed that Slam1 plays a critical role in the transport of TbpB across the outer membrane (Hooda et al 2015). However, the genetic experiments in *N. meningitidis* and the heterologous expression experiments in *E. coli* did not yield a concrete answer as to the exact role of Slam during SLP translocation.

In this study, we developed an *in vitro* functional assay that allowed us to investigate the role of Slam1 in TbpB translocation. Such assays have been previously developed to study the role of outer membrane protein translocons such as the Bam complex (Hagan et al, 2010; Hagan et al, 2011), the autotransporter EspP (Roman-Hernandez et al, 2011), the two-partner secretion system (TPSS) protein B (Norell et al, 2014; Fan et al, 2012) and the lipopolysaccharide translocon LptD (Sherman et al, 2018). By reconstituting Slam1-mediated TbpB translocation *in vitro*, we confirmed that Slam1 acts as an autonomous translocon for the movement of TbpB across the membrane. Furthermore, we discovered that the periplasmic chaperone, Skp, interacts with TbpB and at least another Slam-dependent SLP, named HpuA in the periplasm. We also found that the presence of Skp is crucial for the efficient translocation of TbpB to the surface of *N. meningitidis*. Taken together, we propose a pathway for the localization of surface lipoproteins from the cytoplasm to the surface of Gram-negative bacteria.

## Results

### Purification and incorporation of Slam1 into liposomes

To evaluate the feasibility of characterizing the role of Slam in SLP translocation, we first expressed and tested the function of different homologs of Slam to screen for the most stable Slam-SLP pair for the in vitro study. From our analysis, we found that the Slam1 from *Moraxella catarrhalis* (or Mcat Slam1) expressed well and the purified protein was more stable than other Slam homologs. In addition, co-expression of Mcat Slam1 with Mcat TbpB successfully reconstituted the display of TbpB on the surface of *E. coli* (DE3) cells (Supplementary Fig. 1&2) making Mcat Slam1 a suitable protein for this study. The results also suggested that the components from *E. coli* are sufficient for Slam1-dependent TbpB translocation.

To determine whether Slam1 is an outer membrane translocon working independently from other major translocation system such as the Bam complex, we attempted to reconstitute the Slam1-dependent TbpB translocation with minimal components. First, we tested the incorporation of purified Mcat Slam1-DDM complex into liposomes. Detergent removal allowed for successful insertion of Mcat Slam1 as seen by SDS-PAGE, and western blot analysis using α-His antibodies (Supplementary Fig. 3). To examine liposome insertion, we used the *E. coli* BamABCDE complex as a control. BamABCDE was purified as previously described (Hagan et al 2011) and could potentiate the insertion of the outer membrane protein OmpA into liposomes (Supplementary Fig. 4). Insertion of Mcat Slam1 or Bam complex into liposomes did not affect the stability of liposomes as proteoliposomes containing these proteins were able to float to the top of sucrose gradients upon ultracentrifugation (Supplementary Fig. 5a). Further, to examine the orientation of Mcat Slam1 and Bam complex in proteoliposomes, we incubated the Mcat Slam1 and BamABCDE containing proteoliposomes with proteinase K. The addition of proteinase K led to formation of low-molecular weights bands in an SDS-PAGE gel (marked with asterisk, Supplementary Fig. 5b, left panel) and loss of Slam1 band in an α-His western blot (Supplementary Fig. 5b, right panel), indicating that over 80% of Slam is inserted with its periplasmic domain protruding from the surface - the “inside-out” orientation required for the *in vitro* translocation assay.

### Slam1 proteoliposomes translocate purified unfolded substrate

Once we established a proteoliposome with Slam1 incorporation, we attempted to detect the Slam-mediated transport of SLPs across the bilayer (Hagan et al, 2010) (Fig. 1a). To this end, we purified lipidated functional *M. catarrhalis* TbpB for the assay (Supplementary Fig. 6). Translocation of TbpB was assessed by sensitivity to proteinase K. Only the urea unfolded purified TbpB was successfully translocated into Slam1 proteoliposomes (∼3% insertion), but not in empty liposomes or Bam proteoliposomes (Fig. 1b, 1c). The addition of the Bam complex contributed little to no effects into the TbpB translocation efficiency, suggesting that Bam complex does not involve in this process. Furthermore, the low efficiency of insertion observed for the defined system together with the observation that translocation across the pore only occurs when the TbpB is denatured by urea lead us to hypothesis that there are likely additional periplasmic factors that keep the SLP unfolded for an efficient translocation.

**Figure 1:**
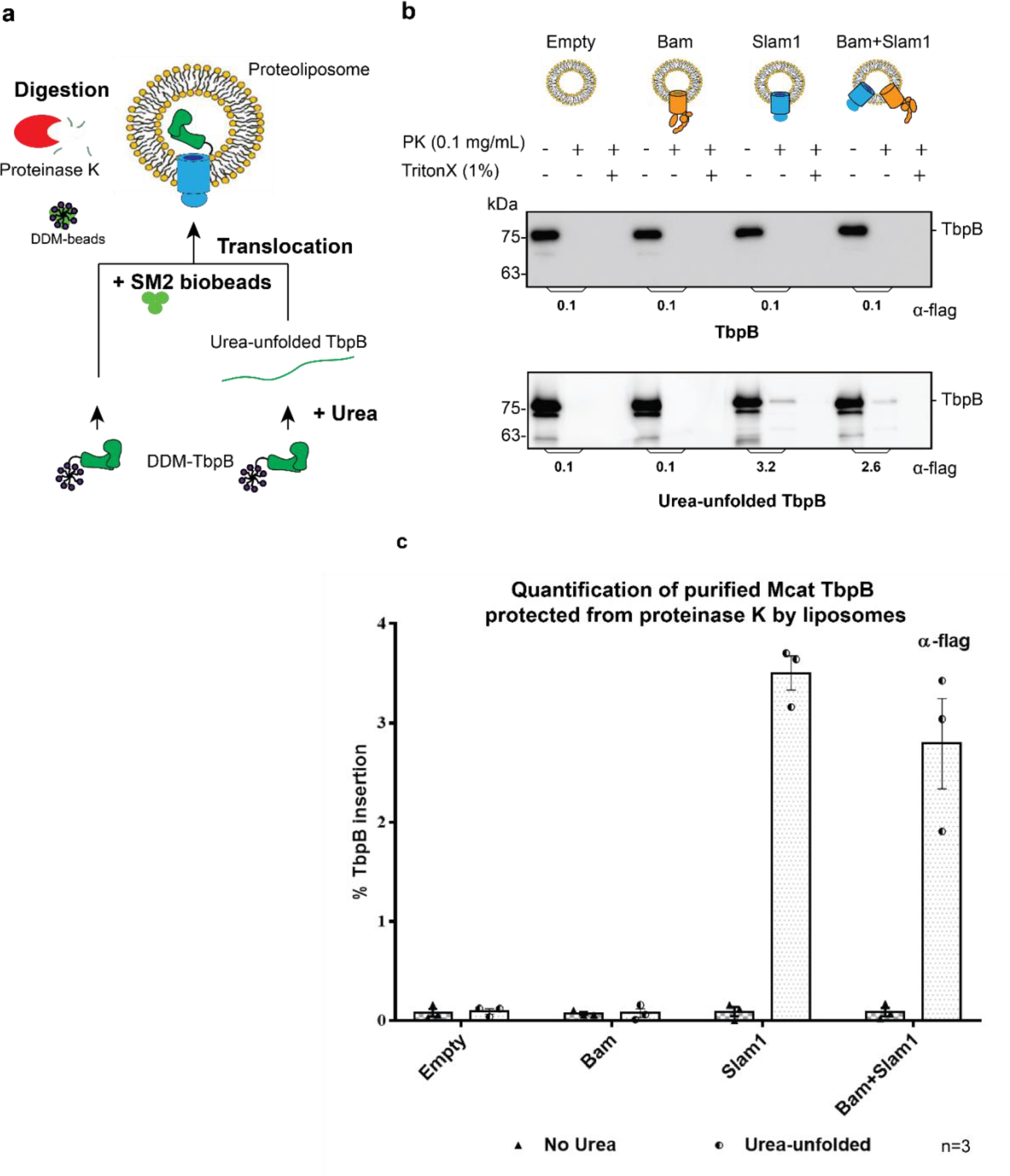
Slam1 is necessary for the translocation of unfolded TbpB. **a)** Model of a defined *in vitro* assay for TbpB translocation*. M. catarrhalis* TbpB (folded and urea-unfolded) is translocated inside Slam1 containing proteoliposomes. Efficiency was calculated based on percentage of TbpB that was protected from proteinase K. **b)** Representative proteinase K protection assay results obtained for Slam1 or Slam1+Bam incubated with purified TbpB (folded or kept unfolded by 8M Urea). Proteoliposomes containing Empty or Bam were used as controls. Each sample was treated with PK or PK + Triton X-100 and examined by western blot. α-TbpB antibody western blots were used to quantify the amount of TbpB. **c)** Quantification of TbpB protection in proteoliposomes through densitometry analysis. The % TbpB protection was calculated by dividing the +PK by input sample. The plot contains results obtained from three biological replicates. Individual data points were included on the graph

### Translocation of TbpB via Slam1 requires periplasmic components but the process is independent from the release of TbpB from the inner membrane

To delve deeper into the mechanism of Slam-mediated SLP translocation and whether additional of periplasmic contents are required for efficient translocation, we examined the translocation of TbpB presented by *E. coli* spheroplasts that lack an intact outer membrane (Norell et al, 2014) (Fig. 2a). TbpB expressed in spheroplasts are displayed on the outer surface of the inner membrane (Hooda et al, 2016). Previous studies have shown that addition of the periplasmic chaperone LolA leads to release of SLPs from spheroplasts in the culture supernatant (Tajima et al, 1998). Hence, we purified *E. coli* LolA and tested LolA-dependent release of Mcat TbpB from spheroplasts (Supplementary Fig.7). Higher amounts of TbpB were detected in the supernatant in the presence of LolA. We incubated the TbpB expressing spheroplasts directly with Slam1 or Bam proteoliposomes and estimated the translocation efficiency of TbpB using a proteinase K assay (spheroplast-dependent translocation). Any TbpB translocated into the lumen of the proteoliposome should be protected from proteinase K digestion. From this assay, we found that proteoliposomes containing Slam1 showed significantly higher protection (40%) compared to Bam proteoliposomes or empty liposomes (5%) (Fig. 2b – upper panel). The protection of TbpB was lost upon the addition of Triton X-100 suggesting that TbpB is shielded from the protease activity by the lipid bilayer of the liposomes. The background protection observed in empty and Bam proteoliposomes is inherent in the procedure, as the assay that was repeated in the absence of any liposomes resulted in similar levels of protection suggesting that the background protection may originate from the spheroplasts themselves. Interestingly, proteoliposomes containing both Bam complex and Slam once again did not improve the efficiency, indicating that the translocation of TbpB does not require Bam complex (Fig. 2c).

**Figure 2:**
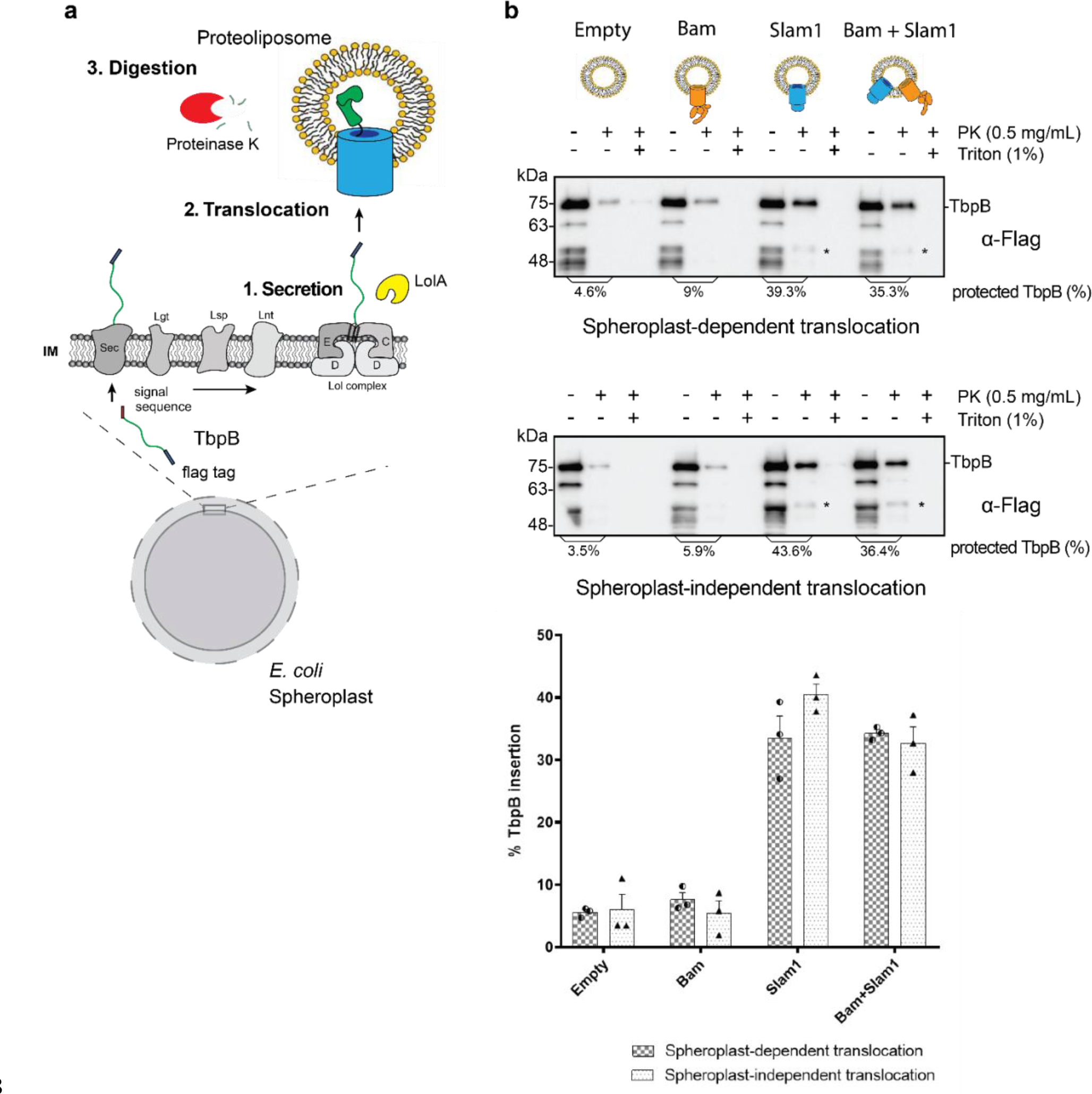
*In vitro* translocation assay for reconstitution of Slam-dependent SLP translocation. **a)** Model of the proposed *in vitro* proteoliposome translocation assay for TbpB secreted directly from *E. coli* spheroplast. **b)** Representative α-flag western blots obtained for the *in vitro* translocation assay. Slam1 proteoliposomes were incubated either with spheroplasts expressing TbpB (spheroplast-dependent translocation, upper panel) or supernatant of spheroplasts that have been induced for TbpB production (spheroplast-independent translocation, lower panel). Empty liposomes and Bam proteoliposomes were used as controls. Proteoliposomes containing Bam + Slam1 were used to test if the Bam complex plays an accessory role to Slam in TbpB translocation. For each proteoliposome, no proteinase K treatment (- PK), proteinase K treatment (+PK) and proteinase K + TritonX-100 treatment (+PK+T) samples are shown. The % TbpB protection shown was calculated by dividing the intensity of the mature TbpB band (∼75 kDa) for each sample by the -PK sample. (*) - partial TbpB fragment which is only seen in the presence of Slam1 proteoliposomes. **c)** Quantification of TbpB protection in proteoliposomes through densitometry analysis. The plot represents data obtained from at least three biological replicates for both spheroplast-dependent translocation and spheroplast-independent assay . Individual data points were included on the graph.

The success of the *in vitro* Slam dependent translocation of spheroplast-released SLPs into liposomes provided an assay to investigate SLP translocation in greater detail. Many outer membrane proteins require inner membrane factors for energy transduction such as TonB-dependent receptors (Pawelek et al, 2006) or chaperone activity TamA (Stubenrauch et al, 2016) to perform their function. Studies of the Lol System have shown that unlike the Lpt system (Sherman et al, 2018), LolA shuttles between the inner membrane and the outer membrane (Szewczyk and Collet, 2016), and hence we predicted that Slam-mediated SLP translocation does not require any inner membrane factors. To validate this hypothesis, we incubated empty, Bam, Slam1 or Bam+Slam1 proteoliposomes with the supernatant isolated from spheroplasts that were expressing TbpB (spheroplast-independent translocation). As seen previously in the spheroplast-dependent translocation assay, we observed TbpB protection from proteinase-K in proteoliposomes containing Slam1 (∼40% protection) and Bam+Slam1 (∼35%) (Fig. 2b - lower panel) but not empty (∼7%) nor Bam (∼5%) proteoliposomes. Interestingly, we did not observe any loss in translocation efficiency between spheroplast-dependent and spheroplast-independent assay (Fig. 2c), confirming that Slam-mediated SLP translocation is independent of SLP release from the inner membrane. This differs from other secretion systems that require partners in the inner membrane who provide energy through ATP/proton motive force (Sherman et al, 2018; Stubenrauch et al, 2016). This finding suggests Slam-dependent SLP translocation is akin to two-partner secretion systems(Fan et al, 2012; Norell et al, 2014; Guérin et al, 2017)

### Periplasmic chaperone Skp interacts with pre-folded TbpB in the periplasm

As previously mentioned above, the Slam1-dependent translocation requires TbpB to be unfolded and hence, we hypothesized that other factors in the periplasm bind the SLPs and prevent their premature folding prior to Slam mediated translocation. To identify periplasmic factors that might be involved in the translocation, periplasmic TbpB complexes were isolated using an affinity flag-tag on its C-terminus. The pulldown fraction was analyzed using mass spectrometry (MS). In this pulldown assay, AfuA – a well-folded periplasmic protein from *Actinobacillus pleuropneumoniae* was used as a negative control to rule out non-specific periplasmic protein interactions (Sit et al, 2015). Skp – a periplasmic chaperone was the only protein that was identified in the pulldown of TbpB but not in the negative control (Table 1). The mass spectrometry results were validated using α-Skp antibody and confirming Skp interacts with TbpB in the periplasm (Fig. 3a). Skp is a homo-trimeric chaperone that binds to unfolded OMPs in the periplasm and is involved in OMP membrane insertion through the Bam complex (Sklar, et al 2007). Our findings suggest that Skp also interacts with TbpB-like SLPs in the periplasm and assists in their translocation across the outer membrane.

**Figure 3.**
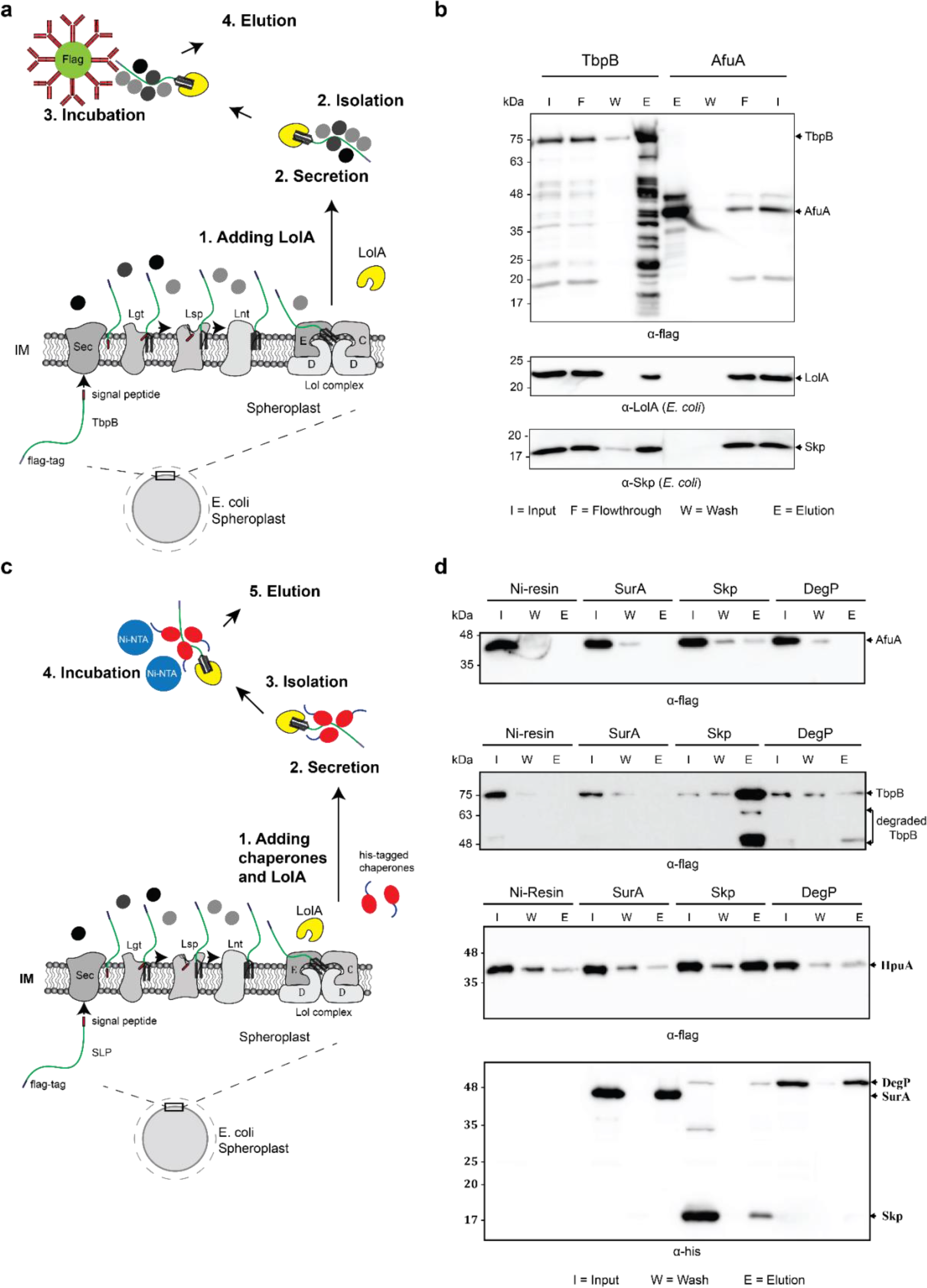
Periplasmic chaperone Skp interacts with surface lipoproteins TbpB and HpuA after being released from the inner membrane. **a)** Model of pulldown assay using the flag-tag on the C-terminus of TbpB. Samples were analyzed using mass spectrometry (summarized in table 1) and examined on western blots. **b)** Representative western blots for TbpB pulldown. LolA and periplasmic chaperone Skp were detected in the TbpB eluted fraction. **c)** Model of reciprocal pulldown assay using the his-tag on the N-terminus of chaperones. Purified his-tagged chaperones (SurA, Skp, DegP) were added to the spheroplast before the induced secretion of SLPs. **d)** Representative western blots of the reciprocal pulldown assay. Only periplasmic chaperone Skp (17kDa) was found to pulldown lipoprotein TbpB and HpuA. Note: Trimeric Skp and DegP have similar molecular weight at 54 kDa.

**Table 1:** Summary of mass spectrometry result. Pulldown samples were left on beads, digested with trypsin and analyzed by mass spectrometry. Data analysis was done by Scaffold 4 software. Cytoplasmic contaminations and proteins that have less than 5 total spectrums count were excluded from the summary table. LolA was detected in both the negative control – AfuA and the protein of interest – TbpB. However, the amount of LolA in TbpB is 5 times higher than amount of LolA in AfuA sample. Skp is the only periplasmic protein that was present in TbpB sample but not AfuA sample, though there were only 9 total spectrum counts. DegP is another periplasmic chaperone that presented in both samples. However, it is known to function as a protease that controls the quantity of over expressed proteins in the periplasm.

|  | AfuA |  |  |  | TbpB |  |  |  |
| --- | --- | --- | --- | --- | --- | --- | --- | --- |
|  | Proteins | Total peptides | % Coverage | Location | Proteins | Total peptides | % Coverage | Location |
| 1 | AfuA | 380 | 78% | P | TbpB | 265 | 31% | - |
| 2 | TufA | 119 | 70% | IM | LolA | 146 | 73% | P |
| 3 | LolA | 36 | 72% | P | TufA | 57 | 59% | IM |
| 4 | OmpF | 12 | 34% | OM | OmpF | 21 | 38% | OM |
| 5 | DegP | 10 | 30% | P | DegP | 15 | 32% | P |
| 6 | FlgH | 12 | 34% | P | OmpA | 15 | 34% | OM |
| 7 | DegQ | 6 | 14% | P | Skp | 9 | 27% | P |
| 8 | OmpA | 5 | 22% | OM | FlgH | 8 | 26% | P |
| 9 |  |  |  |  | DegQ | 5 | 13% | P |

To further validate the interaction between Skp and SLPs, a reciprocal pulldown assay was performed in which a purified His-tagged chaperone was added into the spheroplast prior to the secretion of SLPs. In this assay, we also examined whether Skp interacts with other SLPs such as hemoglobin-haptoglobin utilization protein (HpuA) - a substrate of Slam2 homolog in *N. meningitidis* (Hooda et al, 2016) (Fig. 3b). In addition to Skp, two other periplasmic chaperones which are known to be involved in the transport of OMPs, SurA and DegP, (Sklar et al, 2007) were also examined. The co-immunoprecipitation experiments confirmed that only periplasmic chaperone Skp interacts with the speroplast-released TbpB and HpuA. Chaperone SurA showed no binding to TbpB, while DegP showed modest interaction in line with peptide counts obtained in mass spectrometry (Table 1).

### Periplasmic chaperone Skp is essential for Slam-dependent translocation in *E. coli*

To determine whether Skp is essential for the translocation of SLPs via Slam, TbpB and Slam1 were reconstituted in K12 *E. coli* strains devoid of functional Skp or DegP (as a negative control) (Baba et al, 2006). The presence of TbpB on the surface of *E. coli* was detected using rabbit α-flag antibody, followed by phycoerythrin-conjugated α-rabbit IgG which fluoresces at 575nm. The results showed that only *E. coli* K12 *Δskp* mutant had significant reduction of TbpB’s surface exposure (50%) compared to wildtype cells. Depletion of DegP slightly reduced the translocation of TbpB but this was not statistically significant. No reduction in the expression of either Slam1 or TbpB was observed in western blots. Furthermore, the processing of TbpB by signal peptidase II and subsequence release from the inner membrane was unaffected suggesting the defect in surface display by Skp occurs after the release of TbpB from the inner membrane (Fig. 4a).

**Figure 4:**
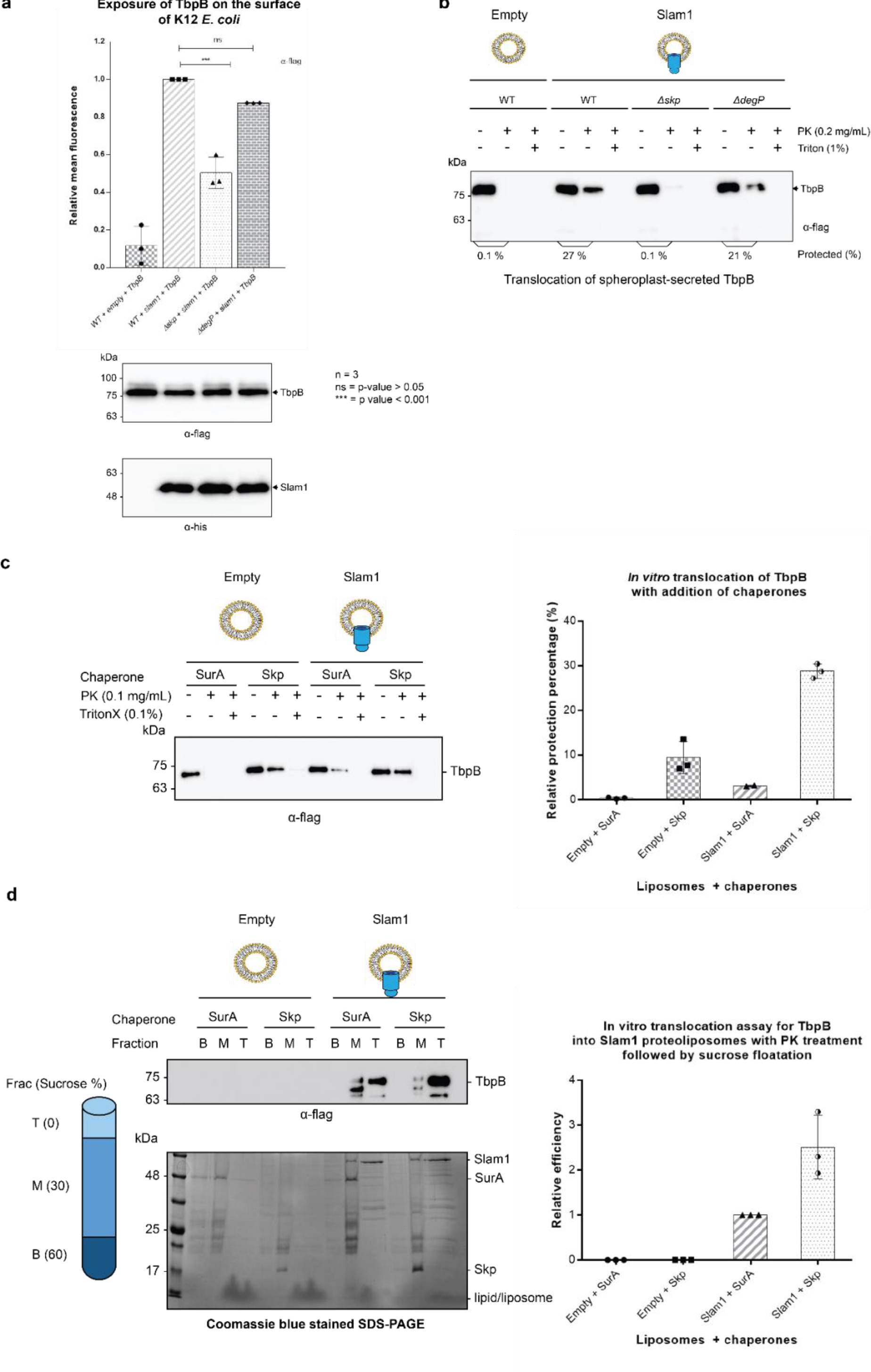
Periplasmic chaperone Skp is required for Slam1-TbpB translocation in the reconstitution systems. **a)** Translocation of TbpB via Slam1 to the surface of *E. coli* K12 mutants. Depletion of Skp significantly reduces the translocation of TbpB to the surface (by 50% - detecting by using α-flag antibody). **b)** Representative western blot of the *in vitro* proteoliposome translocation for TbpB secreted from K12 *E. coli* spheroplast mutants. TbpB secreted from *Δskp* spheroplast fails to translocate into the Slam1 proteoliposomes for protection against proteinase K. **c)** Representative western blot (left panel) and quantification (right panel) of the *in vitro* translocation of purified TbpB into Slam1 proteoliposomes in addition of purified chaperones. Full length lipidated TbpB was unfolded in urea followed by incubation with LolA and either SurA (negative control) or Skp before incubating with empty or Slam1 proteoliposomes and proteinase K digestion. TbpB-Skp complex provided extra protection for TbpB even in the absence of Slam1. **d)** Representative western blot (left panel) and quantification (right panel) of the protected TbpB by the liposomes which were isolated using sucrose flotation assay after proteinase K digestion. Translocation of TbpB into Slam1 proteoliposomes increased by 2.5 folds in the presence of Skp in comparison with the Slam1 proteoliposomes + SurA (positive control). Results are from at least 3 biological replicates. Individual data points were included on the graph.

To further investigate the role of periplasmic chaperone Skp, we leveraged our *in vitro* translocation assay using Slam1 proteoliposomes and spheroplast-secreted TbpB . TbpB that was secreted from K12 spheroplast mutants that lacked Skp or DegP, was incubated with Slam1 proteoliposomes for translocation. The overall results were consistent with the *in vivo* translocation in K12 *E. coli*. In comparison with wildtype- spheroplast TbpB, the *Δskp*-spheroplast-secreted TbpB failed to translocate inside of Slam1 proteoliposomes, while the translocation efficiency of *Δdegp*-spheroplast-secreted TbpB was only marginally reduced (Fig. 4b). This suggests that Slam-mediated translocation of SLPs requires the periplasmic chaperone Skp.

### Skp increases the translocation of purified TbpB in Slam1-containing proteoliposomes

Given that Skp is necessary for Slam-mediated translocation, we hypothesized that addition of purified *E.coli* Skp should increase the translocation efficiency of purified TbpB into Slam1 proteoliposomes. To test this hypothesis, we purified *E. coli* Skp and LolA and added these to urea-denatured TbpB prior to incubation with Slam1-containing proteoliposomes. Periplasmic chaprone SurA was also purified and used as a negative control as we have showed that SurA does not interact with TbpB. As expected, in the presence of Skp, the translocation efficiency of TbpB increased ∼ 5-fold in comparison with SurA (Fig 4c). Curiously, the addition of Skp to the empty liposomes also increased TbpB’s protection by 2 folds. This protection might be from the protease resistance that chaperones provide for their substrates in the periplasm which has previously reported also seen for unfolded OMPs (Yan et al, 2019) (Fig. 4c). Furthermore, this result was consistent with the background protection observed in the spheroplast secretion translocation assays which contained periplasmic components (Fig. 2b). To confirm that the background protection is from the protease resistance of chaperone-substrate complex, the samples were spun down against a sucrose gradient (0-60% w/v) after the proteinase K treatment to isolate the proteoliposomes. The western blots and Coomassie blue stained SDS-PAGE showed a clear separation of the two components (Fig. 4d). While most of the proteins were in the bottom and middle fraction, significant amount of Slam1 and TbpB were found only in the liposomes fraction collected from the top layer. A 3-fold increase in translocation efficiency for TbpB was observed in the presence of Skp compared with SurA and this ratio is consistent with the previous result if accounting for the 2-fold protection coming from the Skp-TbpB interaction. Taken together, these results suggest that Skp potentially plays an important role in the translocation of TbpB to the surface via Slam1, likely through its holdase function.

### Deletion of Skp in B16B6 decreases the exposure of TbpB on the surface of *N. meningitidis*

To examine the role of Skp in the Slam-dependent translocation of SLPs in *N. meningitidis* that contains endogenous TbpB and Slam1, we deleted the gene *skp* (Supplementary Fig. 8, 9) and examined its effect on surface display of TbpB. Such experiments have been previously done in other studies for periplasmic chaperones SurA, Skp and DegQ (homologs of DegP) in *N. meningitidis* in which a single deletion of either one of the chaperones did not affect cell vitality, as well as the expression of OMPs or their insertion into the outer membrane via the Bam complexes (Volokhina et al, 2011). In our study, the deletion of Skp overall did not affect the growth of *N. meningitidis* as the cells reached the optimal OD600 after 12h with a lagging phase at the beginning (Supplementary Fig. 10). In this assay, we used α-TbpB antibody to probe for TbpB on the cell surface. Unlike the two negative controls (*ΔtbpB* and *ΔSlam1*) which completely inhibit the translocation of TbpB, deletion of Skp reduces the amount of TbpB about 50% comparing to the wildtype strain (Fig. 5a – top panel). Interestingly, the expression of either Slam1 or TbpB was not affected which suggests the reduction of TbpB on the surface might be due to the translocation (Fig 5b). This result is consistent with the translocation of Mcat TbpB to the surface of *E. coli Δskp* mutant in which the signal from the C-terminal flag-tag was used to access the surface display of the protein (Fig. 4c). To examine whether these surface exposed TbpB in B16B6 *Δskp* strain is functional, we probed the cells using biotinylated human transferrin (Calmettes et al, 2012). A 5-fold reduction in binding to biotinylated human transferrin was observed for *Δskp N. meningitidis* strain, indicating a significant loss of functional TbpB assembled on the surface of *N. meningitidis* (Fig. 5a – bottom panel). The complementation of Skp from pGCC4 vector successful rescued the translocation of TbpB of the B16B6 Δskp strain back to the wildtype level. Taken together, in the absence of the periplasmic chaperone Skp, less TbpB are translocated to the surface of *N. meningitidis* and these TbpB also fails to be functionally assembled to bind to biotinylated human transferrin.

**Figure 5.**
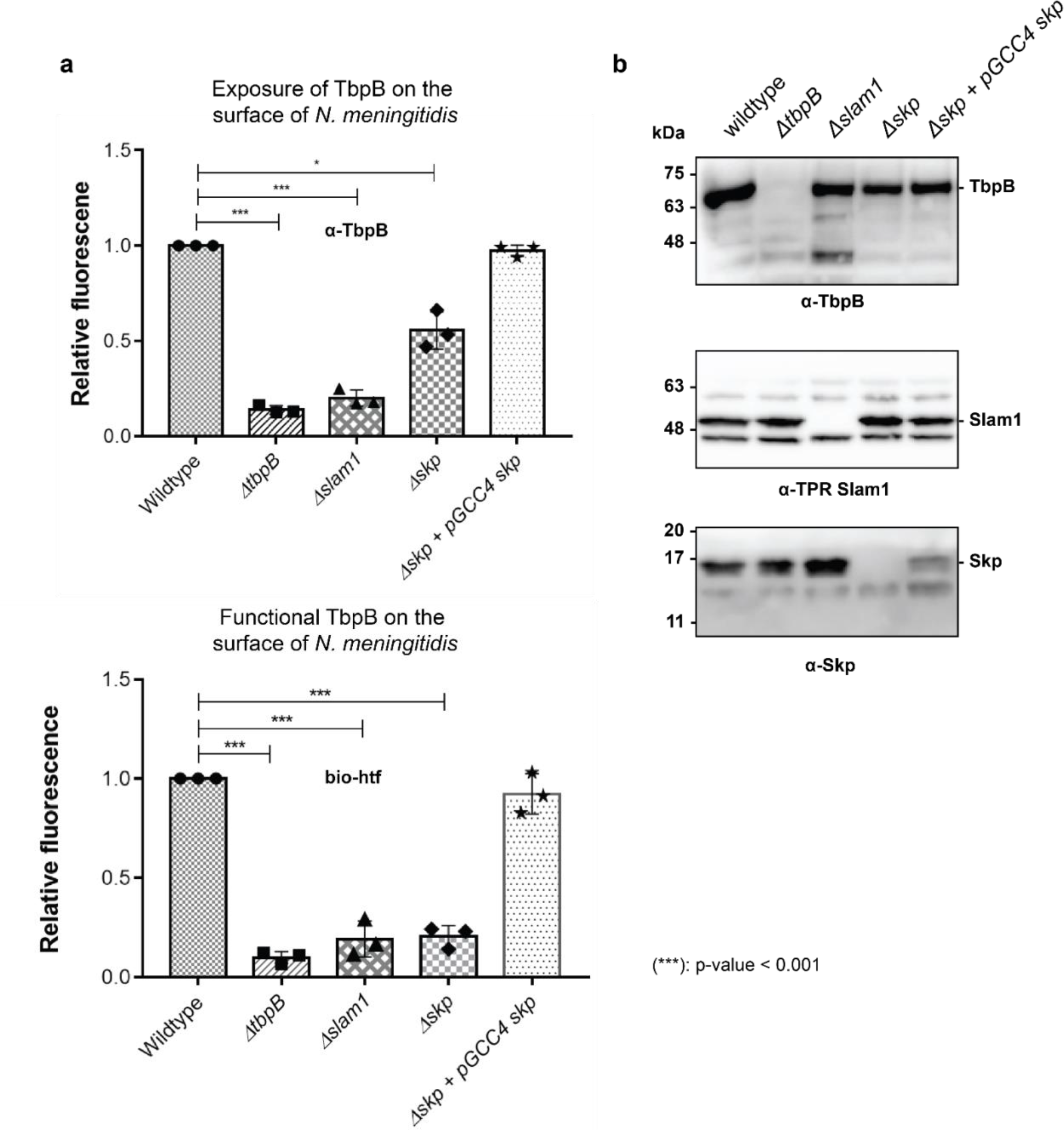
Periplasmic chaperone Skp is important for translocation of TbpB to the surface of *N. meningitidis*. **a)** Relative fluorescence intensity accessing the presence of TbpB (65 kDa) on the surface of *N. meningitidis* mutants using anti-TbpB antibody (exposure - top) and biotinylated human transferrin (functional - bottom). Individual data points were included on the graph. Depletion of *skp* decreased surface exposed TbpB by 50% and the translocated TbpB is non-functional (not binding to biotinylated human transferrin). Complementation of Skp from pGCC4 vector and 1mM IPTG successfully restored the translocation of TbpB and its function. **b)** Representative western blots to access the expression of TbpB, Slam1 and Skp in the *N. meningitidis* strains examined. Depletion of Skp did not affect the expression of OMP Slam1 or TbpB (induced by 0.1mM deferoxamine). Fluorescent assays results are combined from 3 biological replicates and statistically analyzed by one-way ANOVA test.

## Discussion

In this study, we described Slam as an outer membrane translocon responsible for the transport of TbpB-like SLPs to the surface of Gram-negative bacteria. By using an *in vitro* assay to reconstitute the translocation of TbpB across a biological membrane, we showed that Slam1 is necessary to translocate TbpB independently from other outer membrane machineries such as the Bam complex. Unlike other translocons that require energy such as ATP or proton motive force to mediate translocation, we found that Slam1 instead requires periplasmic chaperones to keep pre-folded TbpB available for efficient translocation. In *E. coli* model, Skp was found to interact with TbpB and HpuA in the periplasm after these SLPs were released from the inner membrane. Existing in the trimeric form, chaperone Skp is known to act as a holdase for the pre-folded OMPs as they localize across the periplasm prior to their insertion into the outer membrane by the Bam complex (Sklar et al, 2007; Mas et al, 2019). Given that TbpB and HpuA contain at least one beta-barrel domain similar to OMPs (Calmettes et al, 2012; Wong et alm 2015), Skp might interact with these SLPs in similar manner to keep them in their pre-folded states before translocation (Walton et al, 2009). In our *in vitro* reconstitution assays, the presence of Skp is important for an efficient translocation of TbpB into Slam1 proteoliposomes. The deletion of *skp* in *N. meningitidis* did not affect the expression of neither OMP Slam1 or TbpB but decreased amount of TbpB on the cell surface. Furthermore, the TbpB on the surface of *N. meningitidis Δskp* is not functional as these TbpB fail to bind to biotinylated human transferrin. We do not yet know whether these TbpB were misfolded after the translocation or only part of TbpB was exposed on the surface. Further investigation will be needed to understand how TbpB-Skp complex is recognized and translocated by Slam1, as well as how TbpB is folded once it localized to the surface. Taken together, our data suggest that periplasmic chaperone Skp is required to keep SLPs in their pre-folded states in the periplasm for proper translocation to the surface of Gram-negative bacteria via the Slam translocon.

Combined with our previous work (Hooda et al, 2016), we propose a model for the SLPs localization from the inner membrane to the surface of Gram-negative bacteria (Supplementary Fig. 11). Upon emerging from the Sec translocon, periplasmic chaperone Skp binds to SLPs and keeps them in the pre-folded state. The SLPs are then modified and lipidated before being transferred to the Lol complex. As the SLPs are released into the periplasm, LolA accommodates the N-terminal triacyl lipid group while Skp remains bound to the SLPs to prevent them from prematurely folding prior to being translocated by Slam. Upon their insertion into the inner leaflet of the outer membrane, LolA:SLP:Skp complexes are recognized by Slam for translocation to the cell surface. Drawing from similarities between two-partner secretion (Guérin et al, 2017) and the Slam system, we propose the movement of the SLP across the outer membrane occurs via the Slam membrane domain. Interestingly, Slam substrates such as TbpB or HpuA also contain a lipid anchor which needs to be flipped from the inner leaflet of the outer leaflet, which suggests the presence of a lateral opening in the Slam membrane domain that allows for movement of lipid anchor, similar to ones observed in BamA (Noinaj et al, 2013) and LptD (Gu et al, 2015). Given that Slam translocon seems to require no energy input, we speculate that the folding of SLPs on the surface might provide driving force to further pull the SLPs through Slam barrel domain. High resolution structural and biochemical studies of Slam will be required to reveal the details of SLPs translocation mechanism.

## Materials and Methods

### Bacterial strains and growth conditions

Strains used in this study are summarized in Supplementary Table 1. *E. coli* were grown in LB media containing antibiotics when necessary – 50 μg/mL kanamycin, 50μg/mL erythromycin and 100 μg/mL ampicillin. Cloning procedures were carried out using *E. coli* MM294 competent cells. Protein expression was performed using *E. coli* C43 (DE3) cells for Slam homologs, Bam complex and the translocation experiments (Wagner et al, 2008). *E. coli* BL21 (DE3) cells were used for purification of *E. coli* LolA, SurA, Skp, DegP and B16B6 *N. me* Skp. *In vivo* translocation reconstitution and spheroplast secretion assays were performed using *E. coli* C43 (DE3) or *E. coli* K12 cells from Keio’s collection (Baba et al, 2006). *N. meningitidis* B16B6 strain was used for knock-out study.

### Cloning of Slam, SLPs, LolA and periplasmic chaperones

Genes were cloned into expression vectors by RF cloning (van den Ent and Löwe, 2006) and signal peptides and tags were inserted using round the horn cloning (Liu and Naismith, 2008). pET52 *Nme* HpuA was made by amplifying *hpua* from *N. meningitidis* strain B16B6 and inserting it into an empty pET52b vector. pET52 *Nme* HpuA-flag was made by addition of a flag-tag at the C-terminus of the *hpua* gene in pET52 *Nme* HpuA. pET26 *Ngo* Slam2 construct was obtained by cloning the mature *N. gonorrhoeae* strain MS11 gene *ngfg_00064* and inserting into empty pET26b vector. To be expressed in K12 *E. coli, slam1* and *tbpb* were cloned on pGCC4 and pHERB plasmid respectively. *E. coli lola, sura, skp*, *degp* genes from *E. coli* strain C43 (DE3) genome and *N. meningitidis skp* gene from *N. meningitidis* B16B6 strain were cloned into an empty pET28a vector with an N-terminal 6xHis tag for purification. The constructs used in this study are summarized in Supplementary Table 1.

### Plate reader assay for Slam-SLP *in vivo* translocation assay

Pairs of Slams and SLPs were co-transformed into *E. coli* C43(DE3) or *E. coli* K12 cells. Cells were grown overnight in autoinduction media (Studier, 2005) with appropriate antibiotics as described above. Cells were harvested from the overnight culture by centrifugation at 1500×*g*, 3 mins. Cell pellets were washed gently with PBS + 1mM MgCl2 before incubating with biotinylated human transferrin or rabbit α-Flag antibody (1:200 dilution). After 1-hour incubation, cells were harvested and washed with PBS + 1mM MgCl2. The cells were then incubated with streptavidin-conjugated-phycoerythrin (for biotinylated human transferrin) or α-rabbit IgG-conjugated phycoerythrin (for rabbit-α-flag antibody) with 1:200 ratio for 1 hr. Cells were then harvested, washed and resuspended in PBS + 1mM MgCl2. The samples were aliquoted on a 96-well plate and read on a Synergy 2 (BioTek) plate reader at 488nm and 575nm. OD600 was also recorded for data normalization.

### Purification of Slams

*E. coli* strain C43 (DE3) with pET26 Mcat Slam1 were grown overnight at 37°C in LB + ampicillin. The cells were used to inoculate (1:1000) 6 L of autoinduction media + kanamycin. Cells were grown at 20°C for 48 hours and then harvested by centrifugation at 12200×g for 20 minutes at 4°C. The cell pellets were resuspended in 20 ml/L of 50 mM Tris–HCl pH 8, 200 mM NaCl and cells were lysed using an EmulsiFlex C3 (Avestin). Lysates were spun down at 35000×*g* at 4°C for 10 min. The supernatants were spun down in a 45Ti rotor at 40,000 rpm for 1 hour at 4°C to isolate total membranes. Membrane pellets were homogenized, incubated in 15 ml/L of 50 mM Tris pH 8, 200 mM NaCl, 3% Elugent overnight at 4 °C and the ultracentrifugation step was repeated to remove insoluble membrane pellet. Supernatants containing the soluble membrane proteins were then incubated with 1 ml Ni-NTA agarose O/N at 4°C. Ni-NTA beads were washed three times with 10 column volumes of buffer A (20 mM Tris pH 8, 100 mM NaCl, 0.03% DDM) containing increasing concentration of imidazole. Mcat Slam1 was then eluted in buffer A containing 200 mM imidazole. The protein sample was exchanged into low salt buffer (20 mM Tris pH 8, 20 mM NaCl, 0.03% DDM) using a PD-10 column (GE Healthcare) and then injected onto a MonoQ column (GE Healthcare) equilibrated with low salt buffer. The column was washed with increasing concentration of salt using a high salt buffer (20 mM Tris pH 8, 2M NaCl, 0.03% DDM). Fractions that contained pure Mcat Slam1 were identified using SDS-PAGE gels, pooled, concentrated and stored at -80°C.

For Ngo Slam2 purification, the protocol described above for the expression and purification for Mcat Slam1 was followed up to the NiNTA purification step. Upon elution, Ngo Slam2 samples were concentrated and run on a S-200 column (GE) equilibrated with buffer A. Fractions that contained pure Ngo Slam2 were identified using SDS-PAGE gels, pooled, concentrated and stored at -80°C.

### Purification of Bam complex

The plasmid and protocol for Bam complex purification was adapted from Dr. Bernstein’s group (Roman-Hernandez *et al,* 2014)*. E. coli* strain C43 (DE3) with pJH114 was grown overnight at 37°C in LB + ampicillin. The cells were used to inoculate (1:1000) into 6L of autoinduction media + ampicillin. Cells were grown at 20°C for 48 hours and harvested by centrifugation at 12200×g for 20 minutes at 4°C. Cell pellets were resuspended in 20 ml/L of 50 mM Tris–HCl pH 8, 200 mM NaCl and cells were lysed using an EmulsiFlex C3 (Avestin). Lysates were spun down at 35000×*g* at 4°C for 10 min. The supernatants were spun down in a 45Ti rotor at 40,000 rpm for 1 hour at 4°C to isolate total membranes. Membrane pellets were homogenized, incubated in 15 ml/L of 50 mM Tris pH 8, 200 mM NaCl, 3% Elugent overnight at 4°C, and the ultracentrifugation step was repeated. Supernatants containing the soluble membrane proteins were then incubated with 1 ml Ni-NTA agarose O/N at 4°C. Ni-NTA beads were washed with one column volume with buffer A containing increasing concentration of imidazole. BamABCDE was then eluted in buffer A containing 200 mM imidazole. The protein sample was concentrated and injected onto a S-200 column equilibrated with buffer A. Fractions that contained complete BamABCDE complexes were identified using SDS-PAGE gels, pooled, concentrated and stored at -80°C.

### Liposome and proteoliposome preparation

100 mg of *E. coli* polar lipid extract (Avanti) was resuspended in chloroform (Sigma). The lipid solution was then dried off under N2 gas and resuspended in 10 mL of buffer B (50 mM Tris-HCl pH 7, 200 mM NaCl). The solution was flash frozen and thawed 5 times and stored at -80°C as a 10 mg/mL stock. For each experiment, 1mL of the liposome solution (10 mg/mL) was extruded through a 0.2μm filter (Whatman) to make unilamellar liposomes. The extruded solution was split and the purified outer-membrane proteins (Bam complex and Slam1&2) were diluted 1:5 into the liposome solutions at 1.5μM for Bam and 15μM for Slam1&2. 50 mg of Biobeads SM-2 (BioRad) were added to remove detergent and promote protein insertion into liposomes. Tubes were sealed with parafilm and kept at room temperature with gentle end-to-end rotation for ∼ 2 hours. Beads were changed 2 more time and the proteoliposomes were incubated at 4°C overnight with end-to-end rotation. Proteoliposomes were separated from Biobeads and spun down at 18000×g at 4°C for 5 minutes. The supernatant was kept at 4°C and used for the experiments within a week. The insertion of Slam1&2 and the Bam complex was assessed by SDS-PAGE gels, silver stain and western blots with α-His antibody.

### Sucrose floatation assay

The protocol used for the sucrose floatation assay was adapted from Dr. Müller’s group (Fan *et al,* 2012) with a few modifications. 200 μl of Bam and Slam1 proteoliposomes were resuspended in 1000 μl solution containing 60% sucrose (w/v) and transferred to a 5 ml thin-wall polypropylene Beckman tube. The 60% sucrose was layered with 3.8 ml of 30% sucrose and 200 μl of buffer B. The samples were spun in a Beckman SW 50.2 Ti rotor at 45,000 rpm for 16 hrs at 4 °C. Upon ultracentrifugation, 500 μl fractions were collected from the top. Each fraction was precipitated with 5% TCA, washed 3 times with 100% acetone. The samples were resuspended in 100 μl of 1xSDS buffer and alternate fractions (1^st^, 3^rd^, 5^th^, 7^th^ and 9^th^) were run on an SDS-PAGE gel. Western blots were completed with α-His antibody to estimate the quantity of Mcat Slam1 and BamABCDE present in each of the fractions.

### Purification of periplasmic chaperones from *E. coli* and *N. meningitidis*

Purifications were performed similarly for the soluble proteins*. E. coli* BL21 (DE3) cells expressing either *E. coli* LolA, SurA, Skp, DegP or *N. meningitidis* Skp were grown in 20 mL of LB + kanamycine overnight at 37°C and used for inoculating 2 L of 2YT media. The cells were grown at 37°C to an OD600 ∼ 0.6, induced with 1 mM IPTG and then incubated overnight at 20°C. The cells were harvested the next day by centrifugation at 12200×g for 20 minutes at 4°C. The pellets were resuspended in buffer B (50 mM Tris-HCl pH 7, 200 mM NaCl). Cell lysis was performed using EmulsiFlex C3 (Avestin). The cell lysates were spun down at 35000×g at 4°C for 50 minutes to remove cell debris. Supernatant was filtered through 0.22μm filter and incubated with 1 mL of Ni-NTA beads for 2 hours at 4°C with gentle stirring. The solution was applied to a column and the Ni-NTA beads were subsequently washed 3 times with 10mL buffer B with increasing concentrations of imidazole (10 mM, 20 mM and 40 mM). Proteins were eluted from the Ni-NTA beads by adding buffer B with 200 mM imidazole. The purified proteins were dialyzed overnight in buffer B at 4°C. The proteins were further purified using S75 or S200 gel filtration (GE Healthcare). The purity of proteins was accessed on SDS-PAGE. The proteins were either stored at -80°C or sent for antibody production.

### Spheroplast release assay

The protocol was adapted from Dr. Müller’s group (Fan *et al,* 2012) with a few modifications. Briefly, spheroplasts were obtained from *E. coli* C43(DE3) or *E. coli* K12 cells transformed with either pET52 Mcat TbpB-flag or pET52 *Nme* HpuA-flag or pHERD Mcat TbpB-flag (*E. coli* K12 only). The cells were grown in LB with 100μg/mL ampicillin and induced for expression by 0.5mM IPTG (*E. coli* C43) or 0.1% arabinose (*E. coli* K12) overnight at 20°C. *E. coli* cells were adjusted to have OD600 ∼ 1.0. The cells were harvested by centrifugation at 6800×g for 2 minutes at 4°C. The pellets were then resuspended in 100μL of buffer containing 50 mM Tris-HCl pH 7 and 0.5 M sucrose. The resuspended solutions were kept on ice and converted to spheroplasts by adding 100 μL of buffer containing 0.2 mg/mL lysozyme and 8 mM EDTA with gentle inversion for mixing. The solutions were incubated on ice for at least 20 minutes. The spheroplasts were collected by spinning at 10,000×g for 10 min and resuspended in 100 μL of M9 minimum media containing M9 minimal salts, 2% glucose and 0.25 μM sucrose. Expression of SLPs was resumed by addition of 0.5 mM IPTG or 0.1% arabinose. 10 μM of *E. coli* LolA was added to promote the release of SLPs from the spheroplasts at 37°C. Samples were collected at different time points and spun down at 18000×g for 10 minutes at 4°C to remove spheroplasts. Supernatant at different time points were mixed with SDS loading buffer and run on an SDS-PAGE gel. Western blot analysis using α-Flag antibody to estimate the quantity of TbpB and HpuA released by spheroplasts upon the addition LolA.

### Bam complex functional assay

To test the activity of the Bam complex, the ability of Bam proteoliposomes to potentiate the insertion of spheroplast released OmpA was used. *E. coli* strain C43 (DE3) cells were converted into spheroplasts and recovered in M9 minimal media as previously described. Spheroplasts were then spun down at 18000×g at 4 °C for 10 minutes to isolate the secreted supernatant. Supernatant was spun down again at 60000×g at 4 °C to further remove insoluble and remains of outer membrane. Top 200μL of the soluble fraction was collected and kept on ice. 10μL of iced-cold supernatant was incubated with either 10μL of buffer B (liposome buffer), empty liposome or Bam proteoliposome. Incubations were started every 5 minutes and all reactions was stopped at the same time by adding 5μL of 5×SDS loading buffer. Samples of 0, 5, 10 and 20 minutes were loaded on SDS-PAGE and followed by α-OmpA western blot to access the folding process of *E. coli* OmpA in the presence of Bam proteoliposome.

#### Purification of Mcat TbpB

*E. coli* C43 (DE3) cells was transformed with pET52b Mcat TbpB flag-tag. The cells were grown in 20mL of LB + 100 μg/mL ampicillin overnight at 37°C and were used to inoculate 2L of 2YT + 100 μg/mL ampicillin the next day. Once OD600 reached 0.6, 1mM IPTG was added to induce Mcat TbpB-flag and the protein expression was carried overnight at 20°C. The purification was performed similarly to Slam and Bam outer membrane protein purification protocol. After the membranes were extracted and solubilized in 50mM Tris pH 8, 200mM NaCl and 0.1% DDM, 100μL of flag-beads (sigma) was added into the solution and incubated for 4h at 4°C. The beads were loaded on a gravity column and washed 3 times with 5mL of 50mM Tris pH 8, 200mM NaCl, 0.03% DDM. Mcat TbpB-flag was eluted by adding 500μL of 0.1M glycine, pH 3.5, 0.03% DDM and 100μL of 1M Tris pH 8 was immediately added into the eluted fraction. A280 of the last eluted droplet was measured to determine whether additional volume is needed to elute more protein. All eluted fractions were pooled and concentrated to 0.5 mg/mL. The protein was flash-freezed in liquid nitrogen and stored at -80°C for *in vitro* proteoliposomes assay.

### Dot blot assay for testing function of TbpB

0.5 μl of TbpB (1 mg/ml), TbpA (1 mg/ml), BSA (1 mg/ml) and BamABCDE (1 mg/ml) was spotted on a nitrocellulose membrane. The cells were blocked with 5% skim milk and then developed with a biotinylated human transferrin (50 μg/ml) followed by streptavidin conjugated HRP.

### Translocation assay with purified TbpB

To develop the defined translocation assay, purified TbpB was diluted to 6 μM in buffer B or 8M urea. The TbpB samples were rapidly diluted 1/12 into 50μL of Empty, Bam, Slam1&2 and Bam+Slam1&2 proteoliposomes to bring the final concentration of TbpB to 0.5 μM and urea to 0.66 M. The samples were incubated for 15 min at 37°C with addition of 10mg biobeads. The solutions were isolated and then incubated with proteinase K (0.5 mg/ml) in the presence or absence of Triton X-100 (1%). Samples were incubated at room temperature for 30 min. 5mM PMSF was then added to inhibit proteinase K. Samples were then run on SDS-PAGE, followed by western blotting and α-flag antibody was used to detected TbpB.

### Spheroplast-dependent translocation assay

To develop the spheroplast-dependent translocation assay, we followed the protocol described above for the generation of spheroplasts. Spheroplasts were collected by spinning at 10,000×g for 10 minutes and resuspended in 100 μL of M9 minimum salt media containing M9 minimal salts, 2% glucose, 0.25 μM sucrose, and 10 μM of *E. coli* LolA. Subsequently, 50μL of empty liposomes or Bam, Slam1 or Bam+Slam1 proteoliposomes were added to the separate tubes of the sphereoplasts. Expression of TbpB was induced by the addition of 1 mM IPTG and incubation at 37°C for 15 minutes. Spheroplasts were spun down at 18000×g for 10 minutes at 4°C. Supernatants were collected and treated with the final concentration of 0.5 mg/mL proteinase K in the presence/absence of 1% Triton X-100 and incubated at 37°C for 1 hour. 5mM PMSF was added to inactivate the proteinase K and samples were loaded on SDS-PAGE gels followed by western blots with α-flag antibodies to assess protection from proteinase K activity.

### Spheroplast-independent translocation assay

A similar protocol was performed for spheroplast-independent translocation assay. After 30 minutes of spheroplasts resuming protein expression in M9 media with addition LolA, the solution was spun down at 16000×g for 10 minutes at 4°C. 50 μL of obtained supernatant was incubated with 50μL of Empty, Bam, Slam1 or Bam+Slam1 proteoliposomes for additional 15 mins at 37°C (1:1). The samples were then treated with proteinase K (0.5 mg/ml) in the presence or absence of Triton X-100 (1%) as described in the previous section.

### TbpB pulldown assay

C-terminal flag-tagged TbpB was released from *E. coli* spheroplasts as described above. After 15 minutes of incubating with LolA, spheroplasts were removed by spinning down at 16000×g for 20 minutes at 4°C. 1mL of supernatant was obtained and incubated with 50μL pre-washed flag beads at 4°C for 2h. Beads were spun down at 700×g at 4°C for 10 minutes and supernatant was collected as flow through (FT). Beads were washed 3 times with 1mL of 1× M9 media. Beads samples were sent to mass spectrometry facility (SPARC – Sickkids) for trypsin digestion and analysis. For eluting protein complex, beads were incubated with 200μL of 50mM glycine pH 2.8 at room temperature for 5 minutes. Beads were spun down at 700×g at 4°C for 10 minutes and supernatant was collected as elution (E). All samples were treated with 5× SDS loading buffer and pH was adjusted before loading on SDS-PAGE followed by western blotting. TbpB and AfuA (the negative control) was detected using rabbit α-flag antibody, followed by α-rabbit HPR secondary antibody. LolA was detected using mouse α-his antibody and Skp was detected using mouse α-*E.* coli Skp antibody, followed by α-mouse HPR secondary antibody.

### Chaperone pulldown assays

His-tagged chaperones (SurA, Skp and DegP) were purified as described above. 10μM of each chaperone was added along with 10μM untagged LolA during TbpB/HpuA expression in *E. coli* spheroplasts. After 15 minutes, spheroplasts were removed by spinning down at 16000×g for 20 minutes at 4°C. 1mL of supernatant was obtained and incubated with 20μL pre-washed Ni-resin at 4°C for 2h. Beads were spun down at 700×g at 4°C for 10 minutes and supernatant was collected as flow through (FT). Beads were washed 3 times with 1mL of 50mM Tris 7, 200mM NaCl, 10mM imidazole and 0.1% TritonX-100. Proteins were eluted with 200μL of 50mM Tris 7, 200mM NaCl, 200mM imidazole. All samples were treated with 5× SDS loading buffer and pH was adjusted before loading on SDS-PAGE followed by western blotting. TbpB, HpuA and AfuA (the negative control) were detected using rabbit α-flag antibody, followed by α-rabbit HPR secondary antibody.

#### Reconstitution of Mcat Slam1 and Mcat TbpB in K12 *E. coli* strains (wildtype and mutants)

*E. coli* K12 wildtype, K12 Δ*skp* and K12 Δ*degp* were obtained from the Keio’s collection (Baba et al, 2006). These cells were co-transformed with pGCC4 *mcat slam1* (with N-terminal his-tag) and pHERD *mcat tbpb* (with C-terminal flag-tag). Successfully transformed cells were selected on LB + erythromycin (50 μg/mL) + ampicillin (100μg/mL) plate. Cells were grown in LB media with the appropriate antibiotics until OD600 ∼ 0.6 and then were treated with 0.5 mM IPTG for Slam1 overnight expression. The next day, the cells were spun down at 3,000 rpm for 5 min and the pellets were resuspended in fresh LB media (with appropriate antibiotics), recovered for 30 min at 37°C, 150 rpm. 0.1% arabinose was added into the media to induce the expression for TbpB for 4 hours. The cells were then harvested and ready for plate reader assay with biotinylated human transferrin and α-flag antibody as previous described above

### In *vitro* proteoliposomes translocation with addition of periplasmic chaperones

The assay was modified based on previous assay described above for purified TbpB. In this assay, 10μM of DDM-Mcat TbpB complex was diluted 1:10 in 50mM Tris 7, 200mM NaCl, 8M Urea buffer with addition of 20mg biobeads, 10μM *E. coli* LolA and 30μM *E. coli* Skp or *E. coli* SurA (negative control). The denaturation was performed at 4°C for 2 hours in 1.5 mL microcentrifuge tube with end-to-end rotation. The beads and insoluble were removed by spinning down at 16,000×g for 5 min. 50μL of the supernatant was then incubated with 250μL of either empty liposomes or Slam1 proteoliposomes (to further dilute urea concentration) with 50mg fresh SM2 bio-beads. The solutions were incubated at room temperature for 1 hour and were then treated with 0.1 mg/mL proteinase K or proteinase K + 0.1% Triton-X100 for 15 min. 1mM PMSF was added to inhibit the proteinase K before adding SDS loading buffer for gel electrophoresis and western blot.

For the follow-up sucrose floatation assay, the proteinase K digested proteoliposomes solutions (no TritonX-100 treatment) were mixed with 1mL of 60% sucrose and incubated on ice for 10 min. A layer of 10mL of 30% sucrose was then added on top and incubated on ice for 10 min. 1mL of 50mM Tris 7, 200mM NaCl was used to top up the 13mL polyethylene tube and the solutions were spun at 27,000×g for 18 hours at 4°C using SW45 Ti rotor (Beckman). 1mL of top fraction, 10mL of middle fraction and 2mL of bottom fractions were collected for TCA precipitation (Koontz, 2014). The pellets were resuspended in 1X SDS loading buffer, followed by SDS-PAGE and α-flag western blots to estimate the quantity of Mcat TbpB in each fraction.

### Gene deletion and complementation of Skp in *N. meningitidis*

Restriction free (RF) cloning was used for the following plasmid (Supplementary table 1). To completely replace *skp* gene with a kanamycin cassette, pUC19 Δskp::kan plasmid was cloned to contain the *kan2* gene with upstream and downstream 500bp flanking region of *skp.* The plasmid was used to transform *N. meningitidis* B16B6 strain using spot transformation on BHI plate (Dillard, 2011). The plate was incubated overnight at 37°C, 5% CO2. The lawn within the spot was streaked onto a BHI + 75 μg/mL kanamycin and incubated for 18 hours. Colony PCR was used to select cells that have *skp* deleted and the colony was then grown in 3mL of BHI media + 75 μg/mL kanamycin overnight at 37°C, 5% CO2. The cells were adjusted to have OD600 ∼ 1.0 and 500μL was spun down at 3,000 rpm for 5 min while the remaining cells were used to make 30% glycerol stock and stored at -80°C. The cell pellets were resuspended in PBS buffer. 2X SDS loading buffer was then added for SDS-PAGE, followed by α-*Nme* Skp antibody to confirm the absence of Skp in the B16B6 *Δskp* mutant.

Complementation vector pGCC4 Nme Skp was constructed by cloning the B16B6 *skp* gene into the PacI/FseI site of pGCC4 by RF cloning. The plasmid was used to transform B16B6 *N. meningitidis Δskp* strain using spot transformation. The lawn within the spawn was streaked onto a BHI + 5 µg/mL erythromycin plate and incubated for 36 hours. Colony PCR was used to select cell colonies that have *skp* gene reintroduced. The colonies were then streaked on new BHI (+5 µg/mL erythromycin) with 1mM IPTG plate and incubated overnight. Colonies were collected, resuspended in 1X SDS loading buffer, ran on SDS-PAGE and transferred on PVDF blots. α-Nme skp antibody was used to access the expression of Skp from pGCC4 plasmid in the B16B6 *Δskp* mutant.

### N. meningitidis growth assay

B16B6 *N. meningitidis* wildtype, *Δslam1*, *Δtbpb* and *Δskp* mutant was grown overnight in 2mL BHI +/- 50μg/mL kanamycin. The OD600 was adjusted to 1.0 and 2μL was used to inoculate 200μL of BHI +/- 50μg/mL kanamycin. The cells were grown on a 96 well-plate with 150 rpm shaking at 37°C. The OD600 was recorded every 30 minutes for 24 hours using Nivo microplate reader (VICTOR Nivo).

### Exposure of functional TbpB on the surface of *N. meningitidis* mutants

The cultures were started similarly to the growth assay. After adjusting the OD600 to 1.0, 30μL of cells were used to inoculate 3mL of BHI +/- kanamycin (50μg/mL) in a 15mL culture tube. After 4h, 0.1mM deferoxamine was added to induce expression of TbpB. 1mM IPTG was also added to Δskp + pGCC4 Nme Skp to induce expression of Skp. The cells were grown for 16 hours at 37°C, 5% CO2. Cells were adjusted to have OD600 ∼ 1.0 and 1mL of cells were spun down at 3,000 rpm for 5 min. Cell pellets were washed with 500μL of PBS + 1mM MgCl2 and then resuspended in 200μL of PBS + 1mM MgCl2 + 50μg/mL biotinylated human transferrin (bio-htf) or rabbit α-TbpB antibody (1:200 of unknown concentration) followed by 1h incubation at 25°C. The cells were spun down at 3,000 rpm for 5 min and the pellets were washed 3 times with 200μL of PBS + 1mM MgCl2. The cell pellets were resuspended in 200μL of PBS + 1mM MgCl2 buffer with 50μg/mL streptavidin-conjugated-phycoerythrin (for primary of bio-htf) or 50μg/mL α-rabbit IgG-linked phycoerythrin (for primary of α-TbpB) and incubated for 1 hour. Cells were spun down, pellets were washed 3 times, resuspended in 200μL of PBS + 1mM MgCl2 and transferred into Greiner 96-Well Plates black flat-bottom. Fluorescence intensity was read using microplate reader (Synergy) at wavelength 488nm (excitation) and 575nm (emission). OD600 was measured for normalizing the fluorescent signal.

## Acknowledgments

OmpA antibodies were obtained from Dr. Jan Willem deGier, Stockholm University. Plasmid for expression of the Bam complex (BamA-E) was obtained from Dr. Harris Bernstein at the NIH. Mr. Ashutosh Gupta, Andrew Judd and River Jiang helped in development of the purification protocol for Slams. Funding for this study was obtained from the Canadian Institutes of Health Research (CIHR PJT-148795).

## Author contributions

TFM, SMH and YH designed and conceptualized the study. SMH, YH and RL and did the all the protein purification and liposome experiments. SMH and CCLL cloned the various constructs used in the study. MJ worked on protein purification and structural analysis. SMH, YH, and TFM wrote the manuscript and prepared the figures.

## Materials and Correspondence

All data is available in the main text or the supplementary materials. Correspondence and requests for materials should be addressed to.

## Supplementary Information

**Supplementary Fig. 1.**
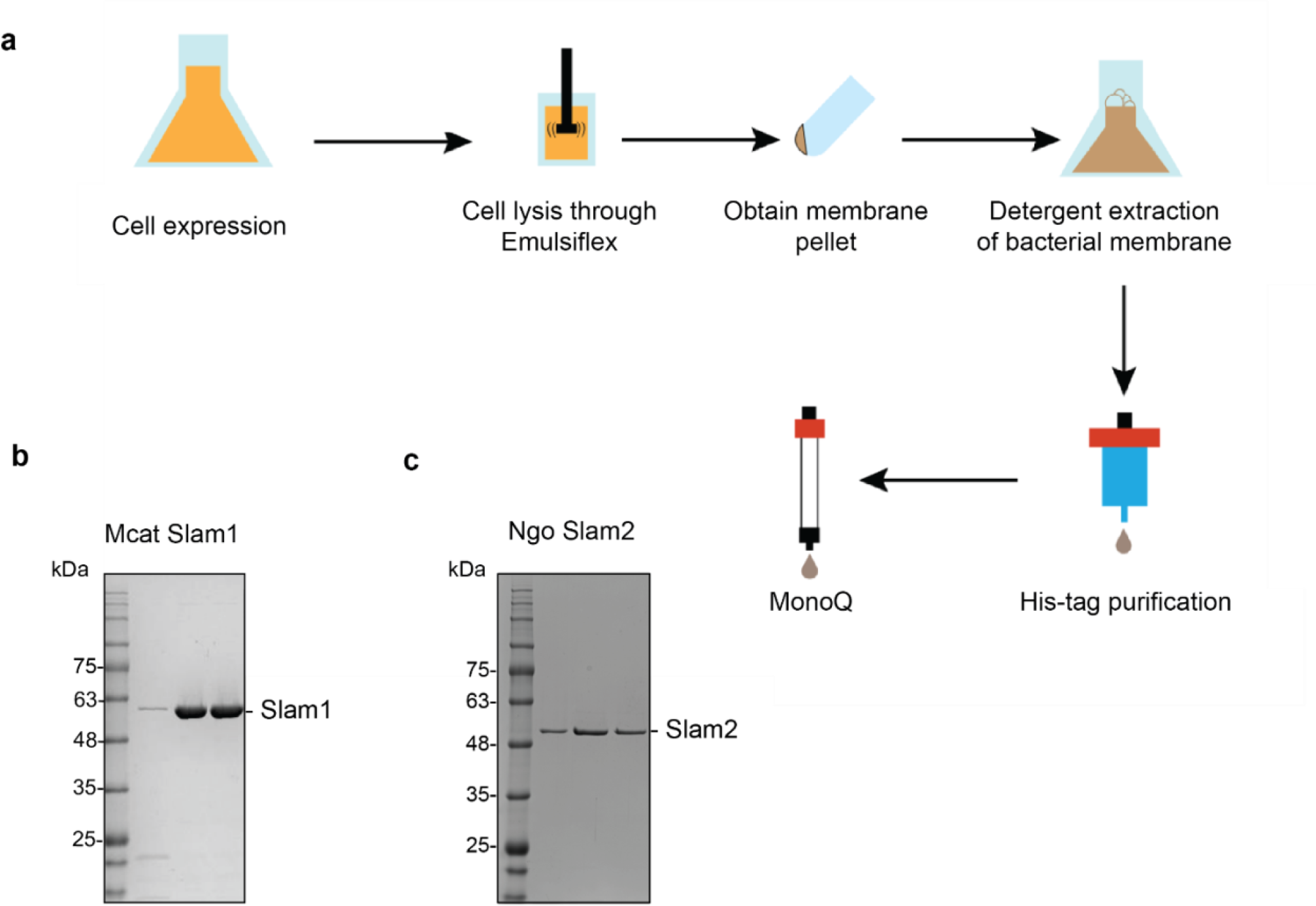
Purification of M. catarrhalis Slam1 and N. gonorrhoeae Slam2. **a)** Overall membrane protein expression and purification pipeline. **b)** SDS- PAGE gels of pure MonoQ fractions from Mcat Slam1 purification. Pure Slam-DDM detergent complex eluted at 60 mM NaCl from a MonoQ column. These were used for the downstream functional assays.

**Supplementary Fig. 2.**
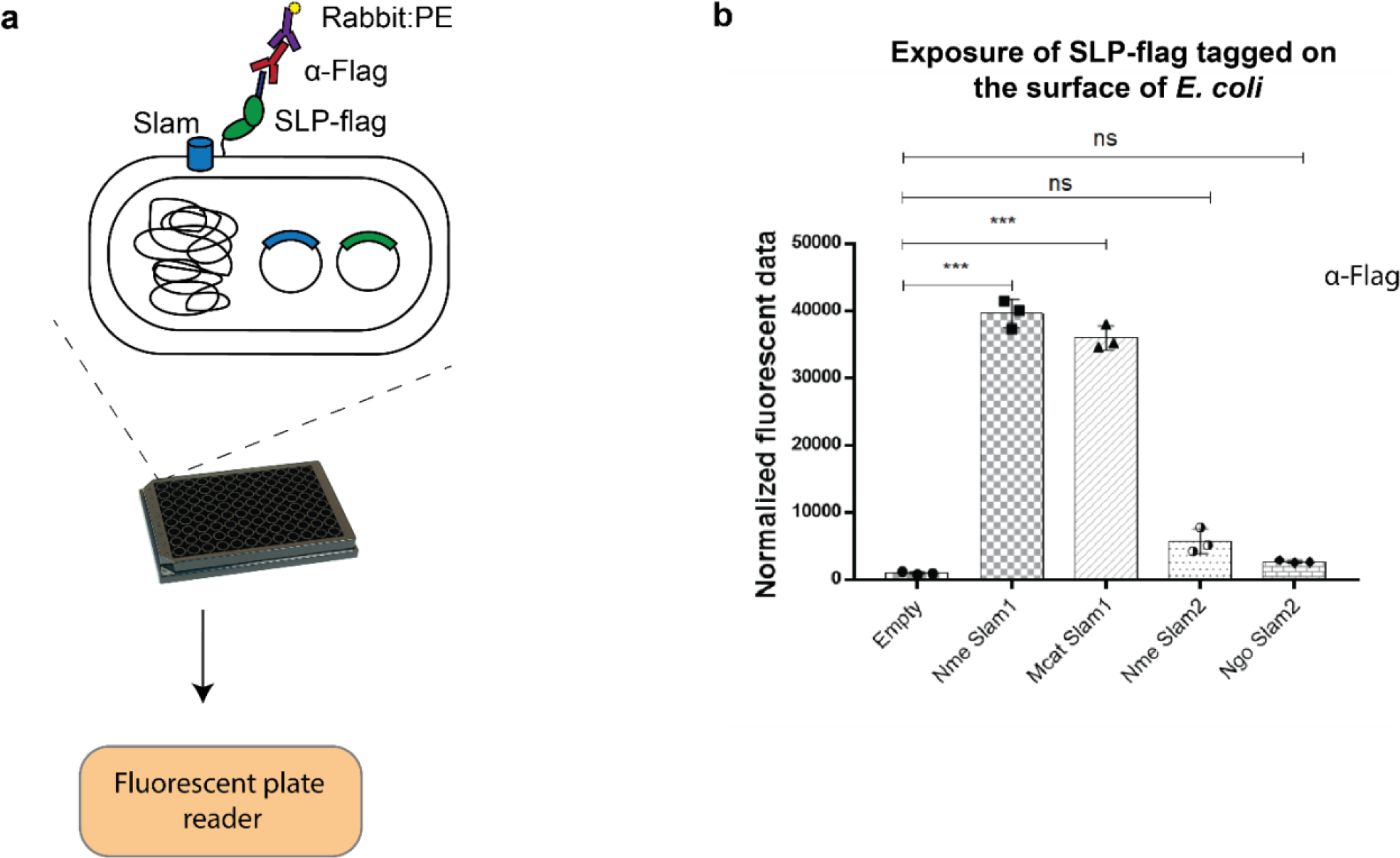
Translocation of Mcat TbpB to the surface of *E. coli* cells. **a)** Plate reader assay was used to examine the function of Slam homologs. Slams and TbpB were co-expressed in *E. coli* strain C43(DE3) and probed with α-Flag antibodies followed by labelling with secondary antibody conjugated with fluorescent probe phycoerthyrin (PE). The fluorescence was quantified using a plate reader. **b)** Quantification of surface display of Mcat TbpB by Slam1 and Slam2 homologs (negative controls). Normalized fluorescence values obtained for each of the Slam homologs is shown. The results represent at least three biological replicates and demonstrate that Mcat Slam1 is functional in translocating TbpB in *E. coli* model. The results also confirm that *E. coli* components are sufficient in reconstituting Slam1-dependent translocation for TbpB.

**Supplementary Fig. 3.**
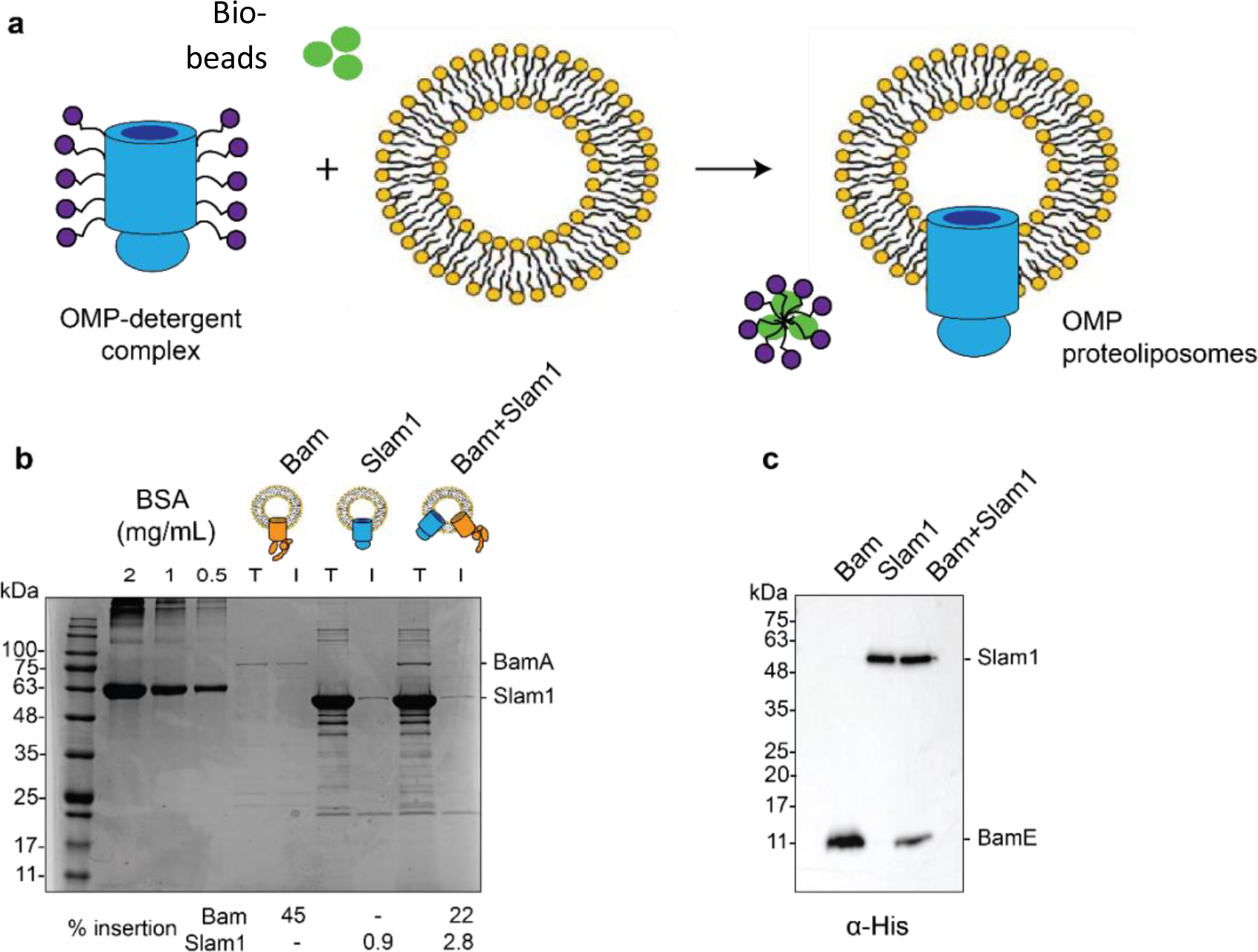
Generation of Slam1 and Bam proteoliposomes. **a)** Protocol used for insertion of outer membrane proteins (OMP) into liposomes. OMP-DDM protein-detergent complexes were diluted (below the critical micellar concentration (CMC) of DDM) into preformed liposomes. Detergent was further removed using SM2 BioBeads. **b)** Quantification of Slam1 and BamABCDE insertion using Coomassie staining. BSA was used as a control for estimating absolute protein quantity. Insertion percentages were calculated by dividing the band intensity of protein inserted in liposomes (I) by total protein incubated with liposomes (T). For the Bam complex, BamA intensity was used to calculate insertion efficiency. **c)** Confirmation of Slam1 and BamABCDE insertion using western blots with α-His antibodies.

**Supplementary Fig. 4.**
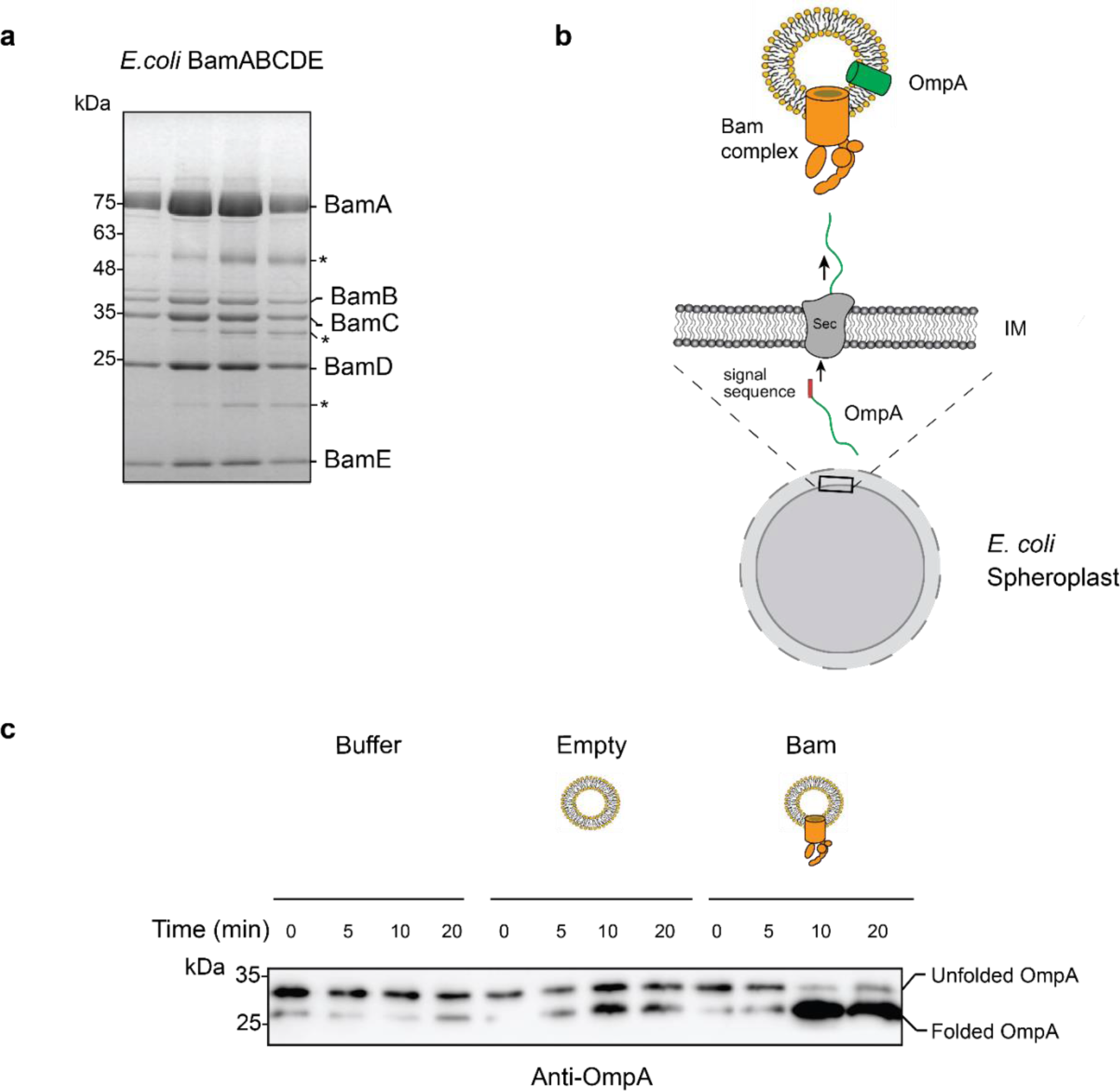
Purification and characterization of E. coli Bam complex. **a)** BamABCDE fractions obtained from a S200 gel filtration column. The BamABCDE complex was obtained using previously described protocols (Ref?). Some non-Bam complex bands (marked in asterisk) were observed and they most likely correspond to common E. coli proteins that have been reported in previous His-tag purified proteins. **b)** Design of an in vitro translocation assay for testing the function of the Bam complex. *E. coli* spheroplasts secrete porins such as OmpA into the supernatant. When incubated with Bam complex proteoliposomes, secreted OmpA is successfully inserted into Bam proteoliposomes. **c)** An α-OmpA western blot to access the folding states of secreted OmpA over time in Tris pH 8 buffer, empty liposome and Bam proteoliposome. Approximately 95% of OmpA achieved folded form in the presence of Bam proteoliposome within the first 10 minutes of incubation while self-folding in empty liposome remained at 50%.

**Supplementary Fig. 5.**
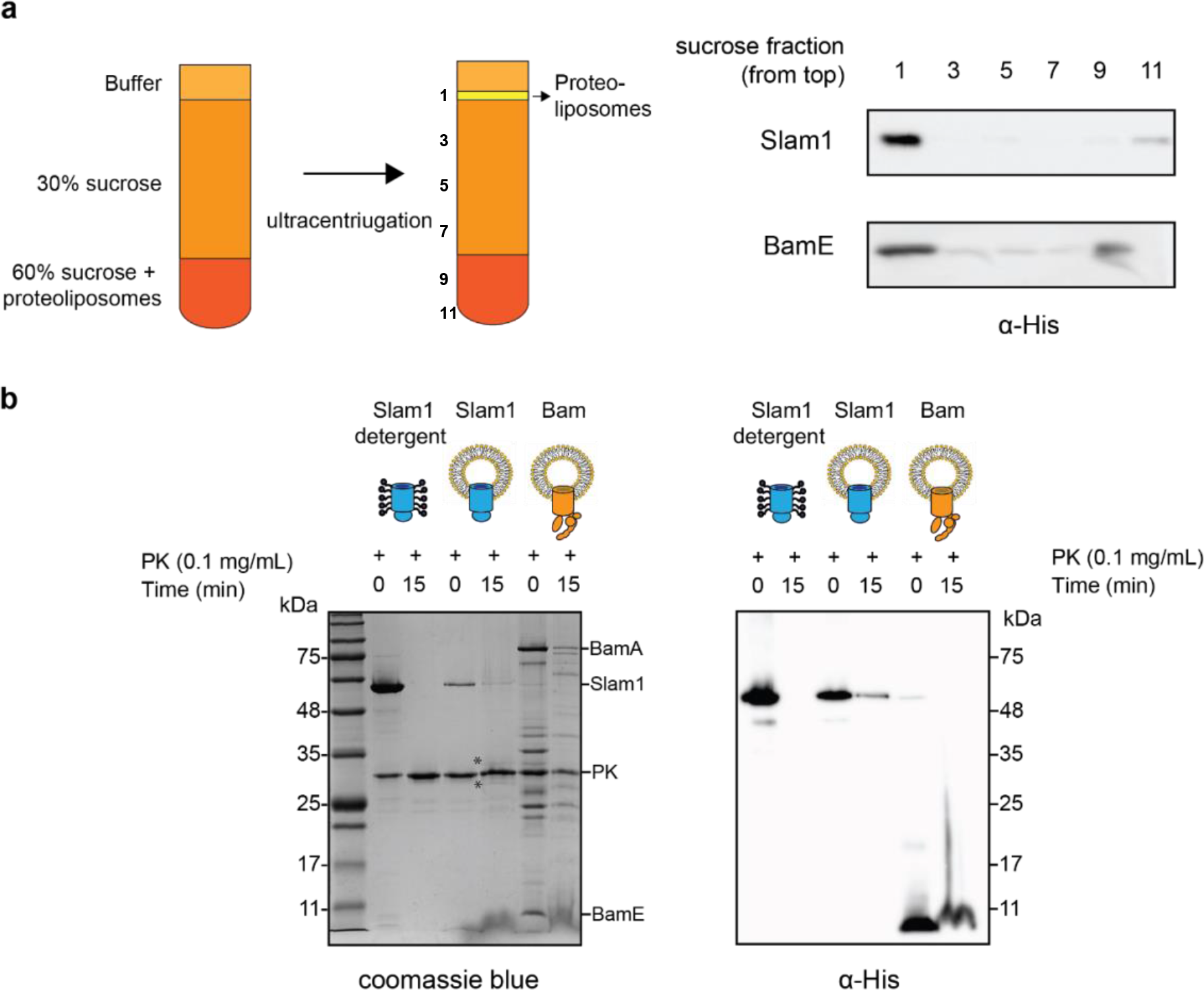
Characterization of Mcat Slam1 and BamABCDE containing proteoliposomes. **a)** Sucrose floatation assay for Slam1 and Bam proteoliposomes. Proteoliposomes were resuspended to a final concentration of 60% sucrose and subsequently layered with 30% sucrose and buffer B. Ten fractions were collected from the top and alternate fractions were run on an SDS-PAGE gel. Western blots using α-His antibodies are shown indicating the amount of protein present in each fraction. **b)** Proteinase K protection assay on Slam1 and Bam proteoliposomes. Proteoliposomes were incubated with 0.1 mg/ml proteinase K for 20 minutes at room temperature. Coomassie blue stained gel and α-His western blot were used to access orientation of the proteins in liposomes. Approxmately 18% of Slam1 inserted with N-terminal his-tag residing in the lumen of liposomes and was protected from PK digestion. Percentage protection was calculated using densitometry. Asterisk (*) indicates fragments of Slam1 (potentially C-terminal barrel) remaining after PK shaving.

**Supplementary Fig. 6.**
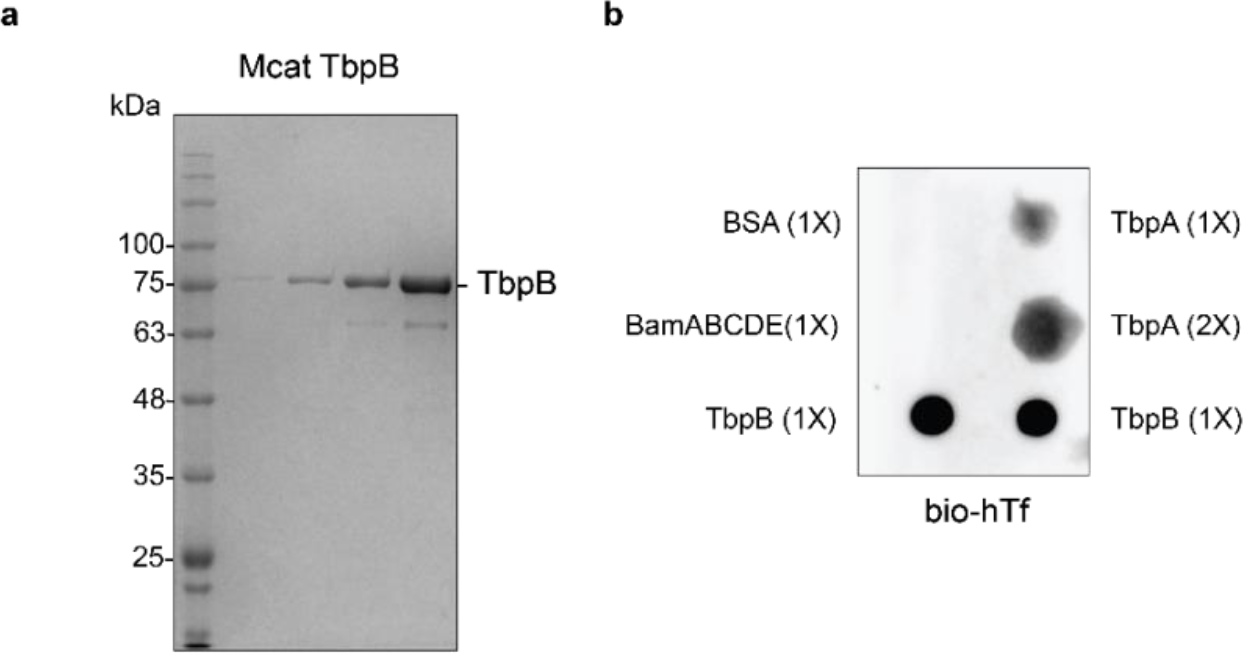
Purification and functional characterization of *M. catarrhalis* TbpB. **a)** Purified TbpB-DDM complexes obtained from a S200 size chromatography column. The sample was subsequently used for the *in vitro* translocation assay. **b)** Dot blot assay with biotinylated human transferrin (bio-hTf) for detecting the function of TbpB. 0.5 μl TbpB and respective controls (TbpA, BamABCDE and BSA) were spotted on nitrocellulose membrane and blotted with bio-hTf (50 μg/ml) followed by streptavidin-HRP. TbpB bound tightly with bio-hTf indicating it is functional.

**Supplementary Fig. 7.**
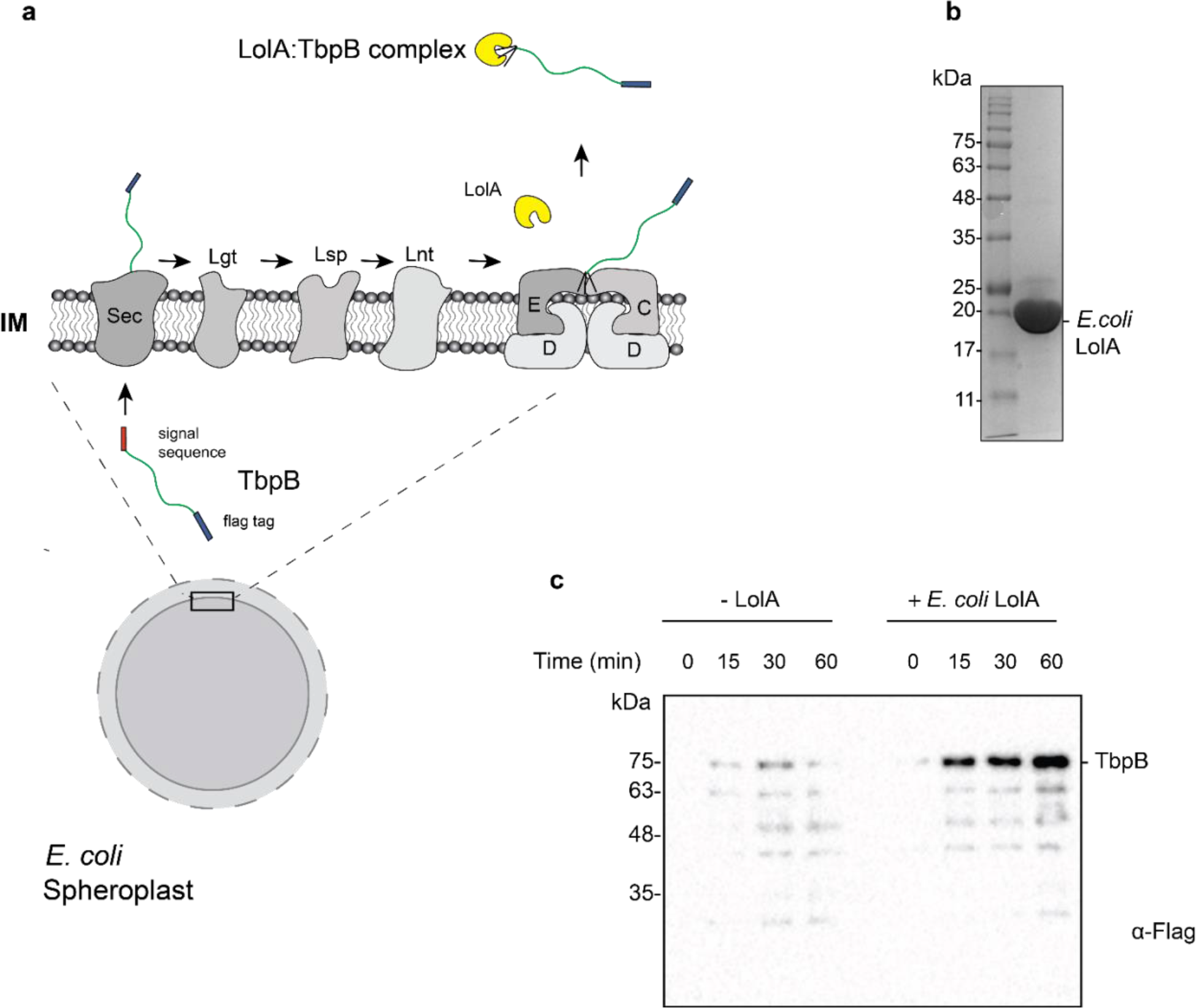
Purification and functional characterization of LolA from *E. coli*. **a)** Model of the release of SLPs from spheroplasts upon addition of purified *E. coli* LolA. SLPs are synthesized in the cytoplasm and transported to the periplasm via the Sec translocon. After the addition of the lipid anchor, SLPs are transferred to the LolCDE complex and released into the periplasm by LolA. In spheroplasts, SLPs are released into the supernatant upon LolA addition. **b)** Uncut 6x-His tagged *E. coli* LolA (∼22 kDa) after Ni-NTA affinity chromatography purification. **c)** Release of Mcat TbpB in the presence of purified LolA over time course of 60 minutes. *E. coli* cell pellets were converted into spheroplasts and induced for expression of TbpB in the presence or absence of 10μM LolA. Samples were collected every 15 minutes, spun down at 13,500 rpm for 5 mins and the supernatants were loaded on SDS-PAGE. The amount of TbpB (∼ 75 kDa) that was released into the supernatant in the presence and absence of LolA was accessed using a α-Flag antibody western blot. Lower bands are degradation product of TbpB which also increased over time.

**Supplementary Fig. 8.**
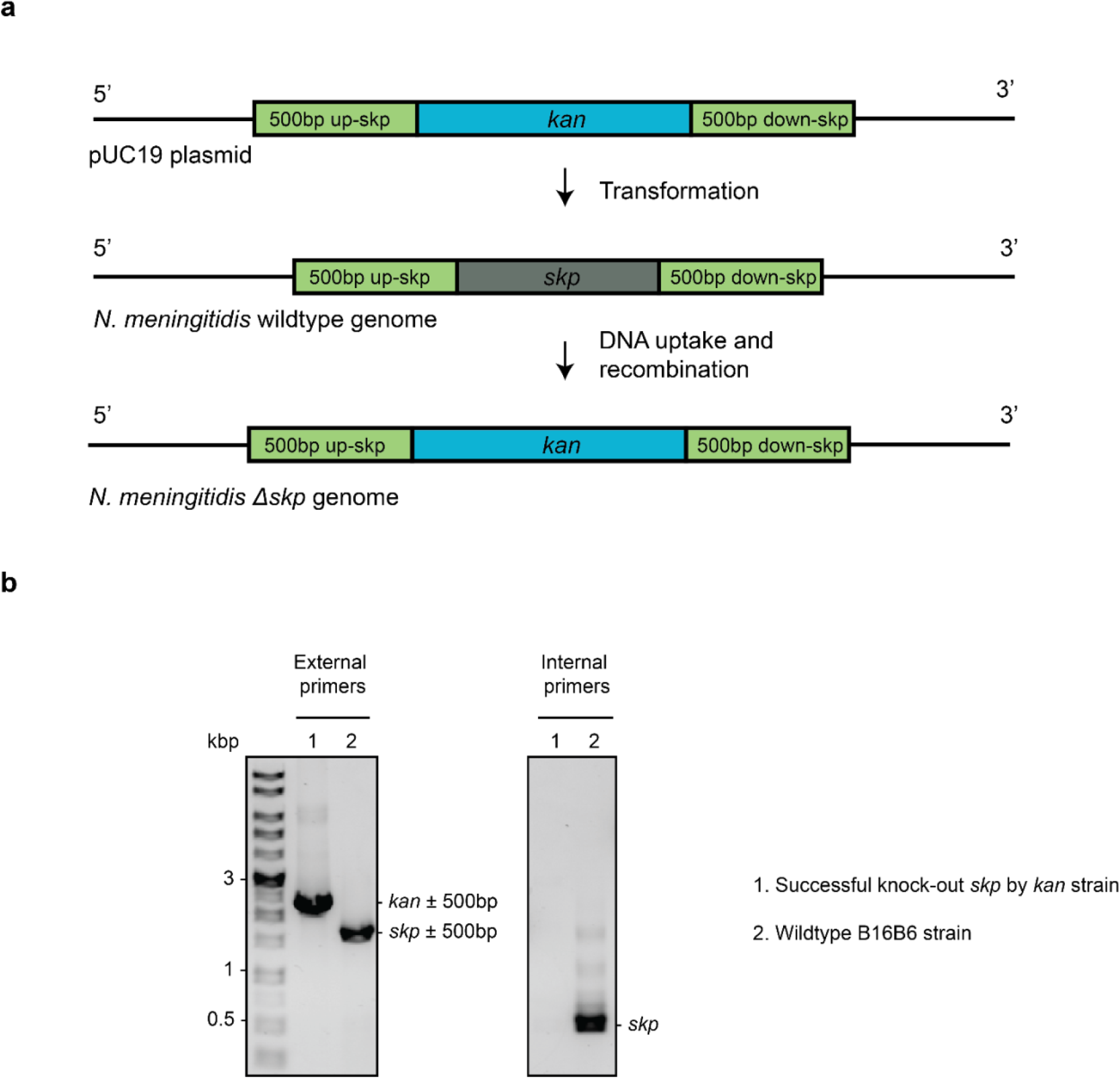
Deletion of Skp in *N. meningitidis.* **a)** The complete knock-out process using spot transformation. **b)** Agarose gel of single colony PCR using external primers (target 500bp up/down-*skp*) and internal primers (target internal of *skp* gene) to confirm the successful replacement of *kan* for *skp gene*.

**Supplementary Fig. 9.**
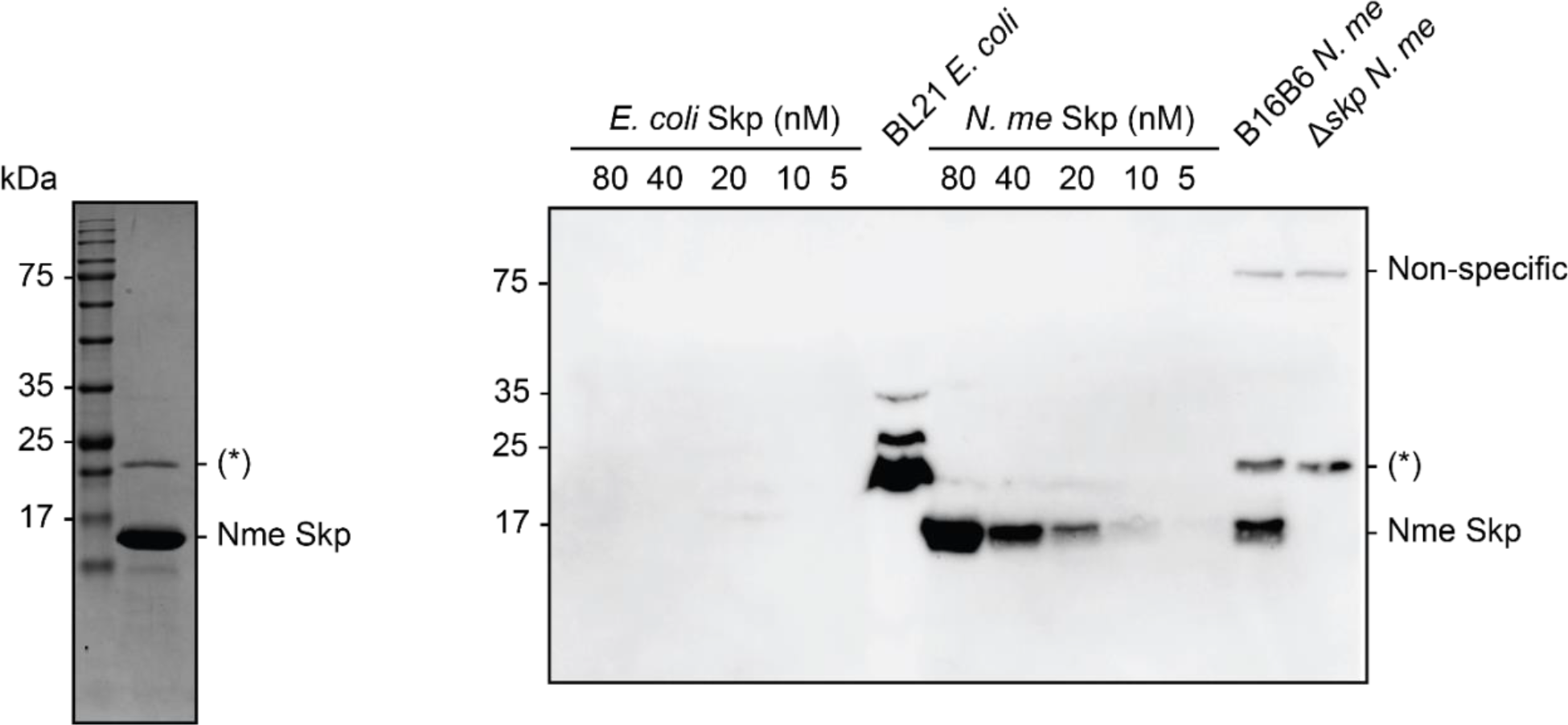
Purification of Nme Skp for antibody production and antibody test. Coomassie blue stained SDS-PAGE of *Nme* Skp after S75 gel filtration to access its purity prior to antibody production (left pannel) and α-Nme Skp antibody western blot to validate the antibody and confirm the knock-out of Skp (17kDa) in B16B6 *N. meningitidis* (right pannel). (*) is the contamination from BL21 *E. coli*. There is no cross-reactivity of *Nme* Skp antibody with *E. coli* Skp.

**Supplementary Fig. 10.**
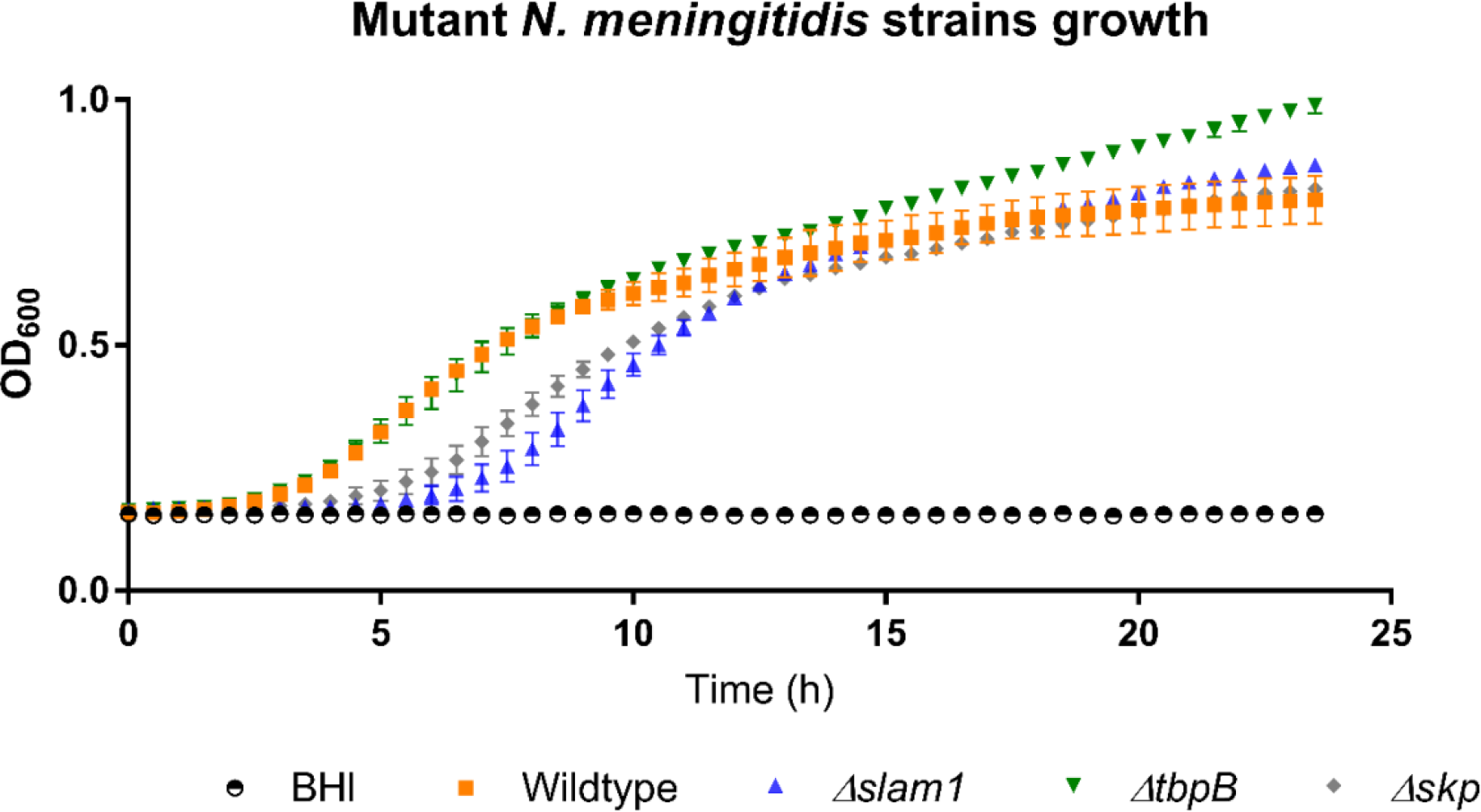
Growth of B16B6 *N. meningitidis* strains in BHI media. OD600 was recorded every 30 minutes over 24 hours period. *Δskp* and *Δslam1* mutants were lagging behind but reached OD600 of 0.7 eventually.

**Supplementary Fig. 11.**
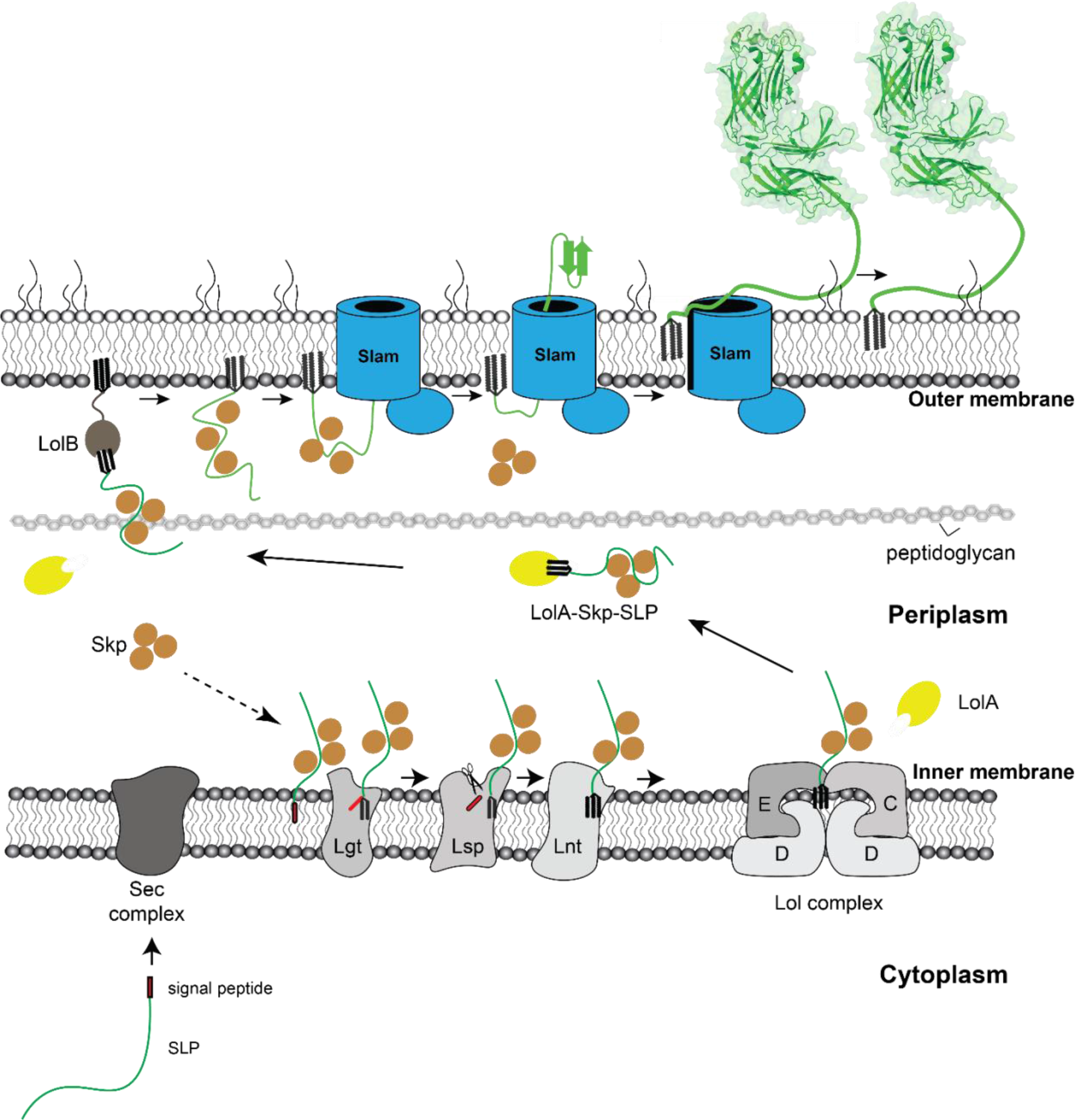
Proposed model of localization for Slam-dependent surface lipoproteins in Gram-negative bacteria. Once the surface lipoproteins emerged from the Sec complex, periplasmic chaperone Skp binds to the SLPs to prevent early folding while their N-terminus is modified and lipidated before getting passed to the Lol complex. LolA then binds to the lipidated SLPs and releases the proteins to the periplsm while Skp stays bound to prevent SLP folding. LolB in the OM serves as the receiver for the LolA-SLP-Skp complex and then inserts the lipidated SLP-Skp complex into the inner leaflet of the OM. The “specificity motif” present at the C-terminus of the unfolded SLP is identified by Slam and then transported across the outer membrane through the Slam channel and folds rapidly. The folding of SLP on the other side triggers the release of chaperone Skp, allowing Slam to pull the rest of the protein across the OM. Once the entire length of the SLP is transported, the Slam lateral gate allows the lipid anchor to “flip” from the inner leaflet to the outer leaflet of the outer membrane.

---

**Supplementary Table S1:**
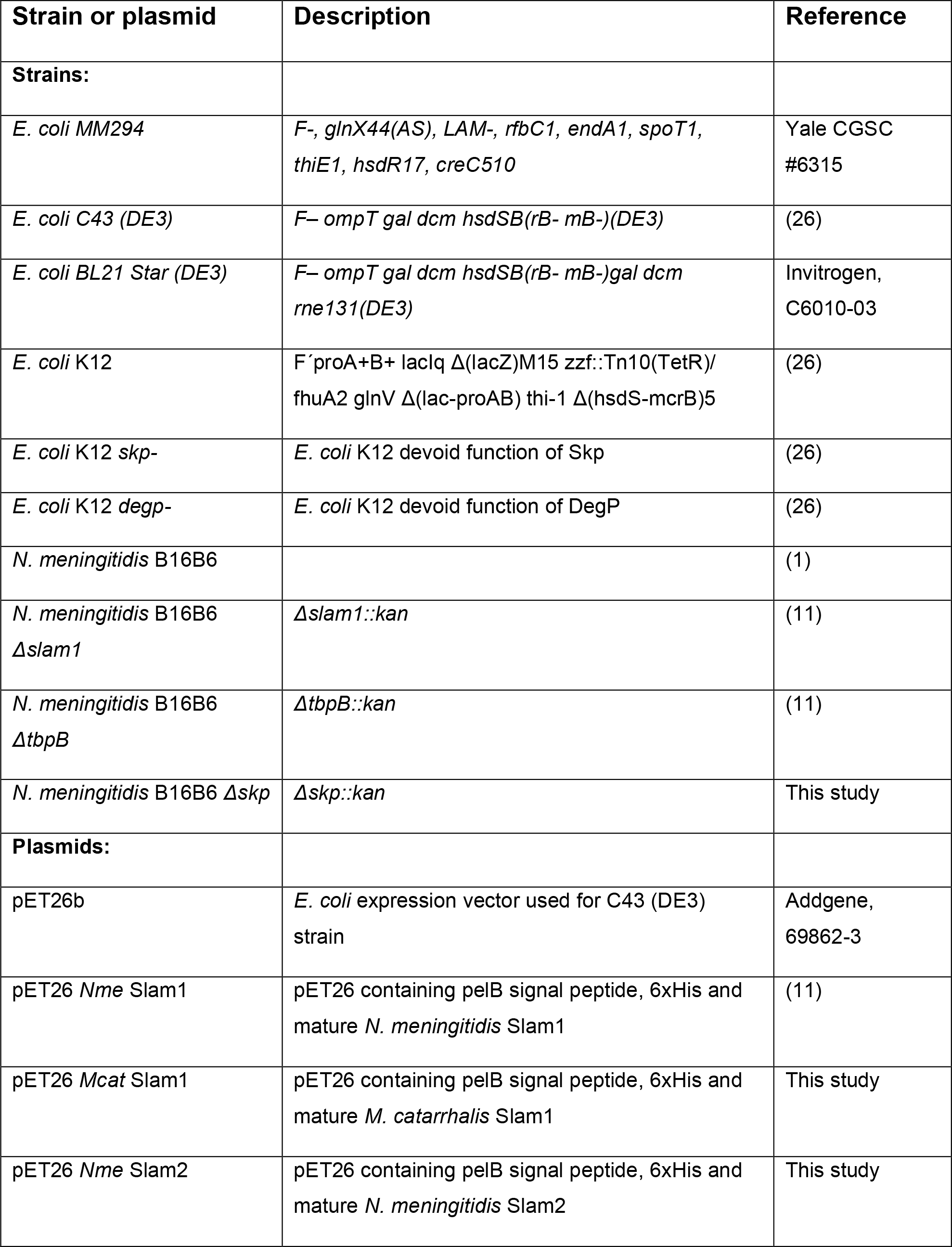
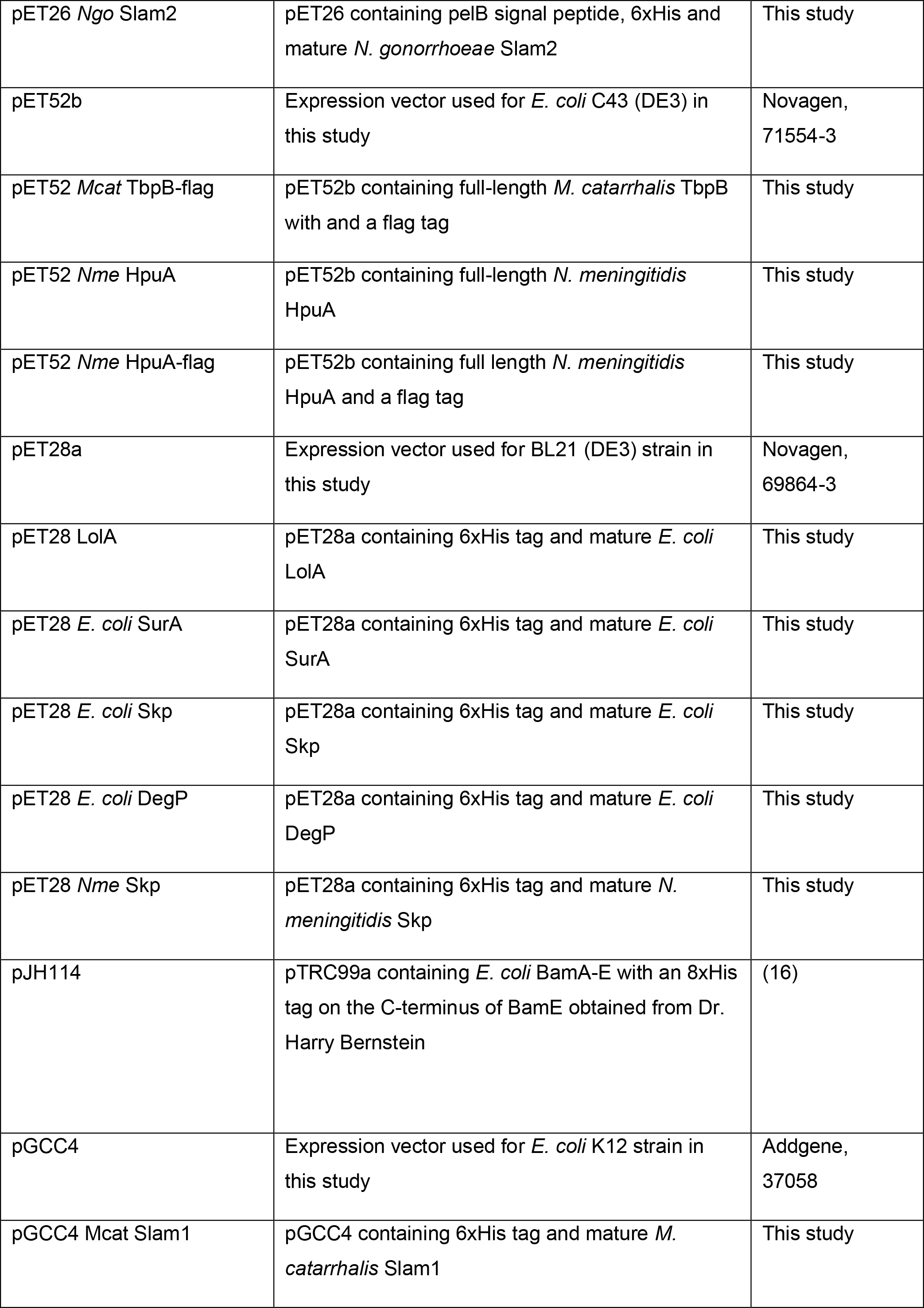
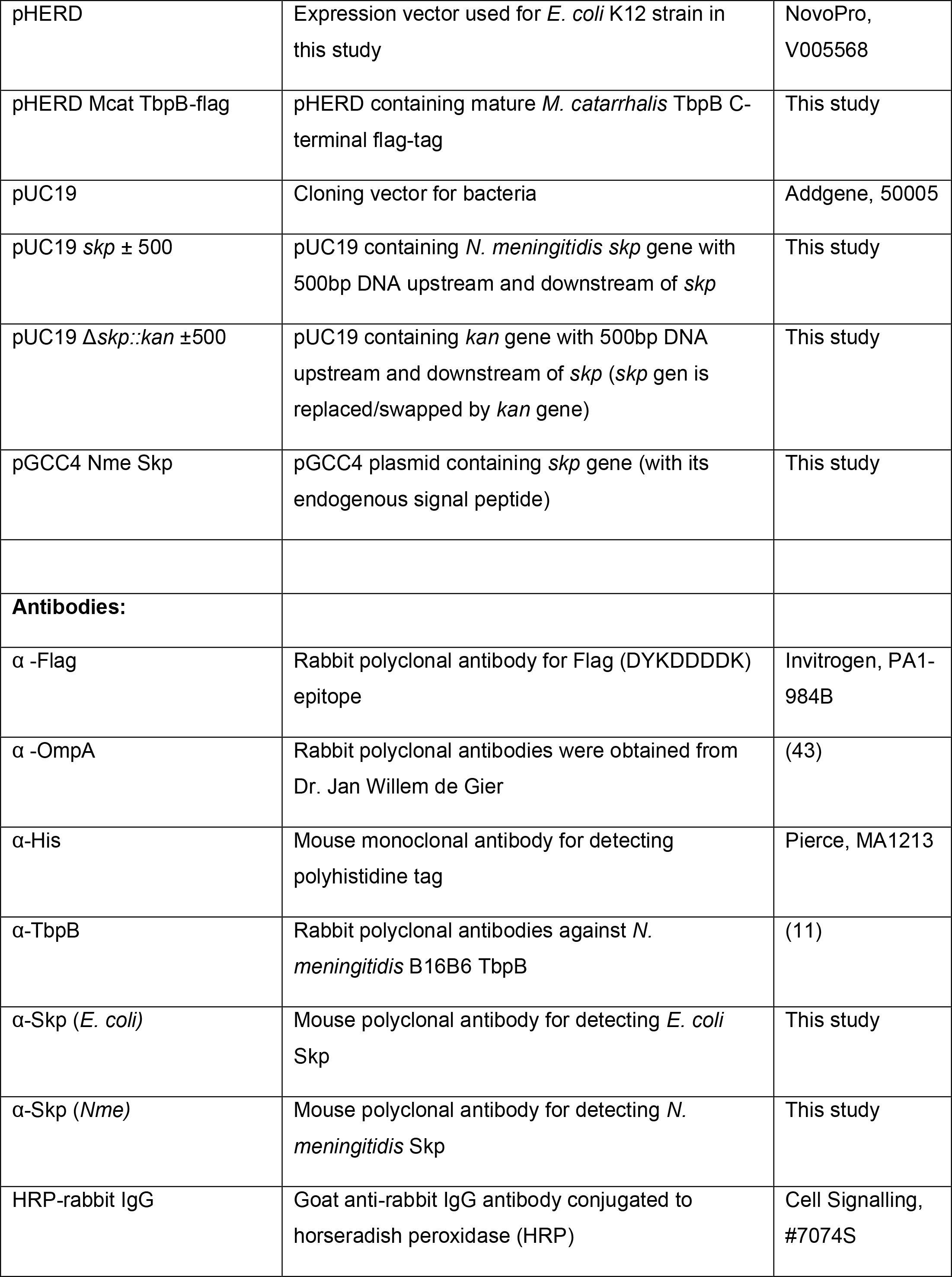
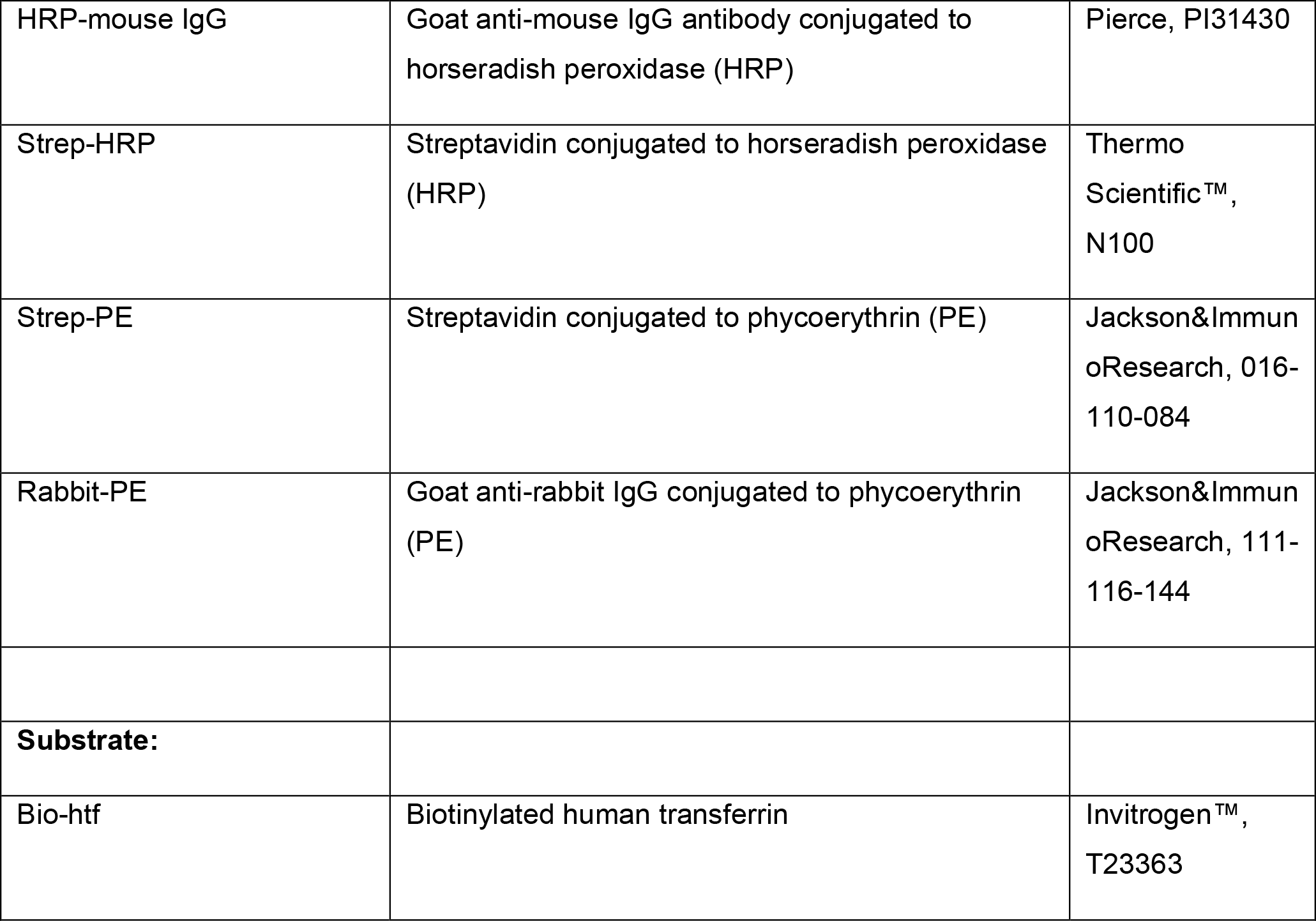
Strains, plasmids and antibodies used in this study.

